# Systematic assessment of the biological impact of cellular deconvolution on downstream analyses of disease transcriptomes

**DOI:** 10.64898/2026.08.05.742993

**Authors:** Sanga Mitra, Maziya Ibrahim, Manikandan Narayanan

## Abstract

**Background:** Cellular deconvolution methods estimate cell-type proportions from bulk RNA-seq data, typically using single-cell RNA-seq–derived signatures, enabling separation of disease-associated transcriptional changes into composition-driven and cell-intrinsic effects. However, these approaches depend on model assumptions and the stability of cell-type signatures, and it remains unclear how deconvolution-related uncertainties influence downstream analyses and biological conclusions.

**Results:** We systematically evaluated the effect of cell-type correction on disease-relevant transcriptomic insights, using Alzheimer’s disease (AD) as a model and the Mount Sinai Brain Bank cohort as a primary dataset. Applying dtangle, selected after comparison with another deconvolution approach, we estimated cell-type proportions across four brain regions and assessed how correction reshaped differential gene expression and pathway enrichment. Cell-type correction (CTC) markedly altered differentially expressed gene (DEG) profiles in a region-dependent manner: the superior temporal gyrus lost all significant signals, while the frontal pole gained DEGs with improved cross-region concordance. At the pathway level, correction shifted enrichment from synaptic loss and immune activation toward suppression of stress-response and immune regulatory programs, suggesting that composition changes partly obscure cell-intrinsic regulatory signals. Overlap with AD genome-wide association study loci and replication in an independent cohort indicated that cell-intrinsic changes are more consistently validated than composition-driven changes. Notably, *KCNN2* and *RIMS1*, not currently recognized as canonical AD biomarkers, emerged as robust transcriptional signatures, potentially reflecting both composition-driven and cell-intrinsic dysregulation and warranting further investigation.

**Conclusions:** Parallel evaluation of uncorrected and CTC analyses distinguishes composition-driven from cell-intrinsic transcriptional effects and highlights robust disease signatures in heterogeneous tissues such as the brain.

## Introduction

Cellular heterogeneity is a defining characteristic of complex tissues such as the brain, where transcriptional changes during disease progression arise from both intrinsic regulatory alterations within cells and shifts in cellular composition (i.e., proportions of different cell types within the tissue). Although single-cell and spatial transcriptomics measurements can disentangle such macro (between cell-types) and micro (within cell-types)-heterogeneity effects [1], they are not scalable to very large cohorts due to cost and resource constraints. In addition, most data that currently exist from large cohorts are derived from bulk RNA-seq, where composition-driven artifacts can obscure true disease-relevant signals. Substantial differences in cell-type frequencies can also persist across individuals. Further single-cell profiling cannot be performed on the original samples, limiting analyses to existing bulk RNA-seq data.

To address this challenge, cellular deconvolution methods have been developed to estimate relative cell-type proportions from bulk transcriptomic data mostly using reference cell-type signatures derived from single-cell datasets [2,3]. Many such methods have been proposed (e.g., CIBERSORT [4], dtangle [5]; and several others cited in [6–8]), and many studies have used deconvolved bulk data to identify conserved molecular signatures across diverse tissues [9]. In many applications, the estimated cell-type proportions are subsequently incorporated as covariates in downstream analyses—a procedure commonly referred to as cell-type correction (CTC)—with the goal of mitigating composition-driven confounding in differential expression analyses.

Despite the widespread use of deconvolution methods, an important question remains insufficiently explored: how does cellular deconvolution influence the biological conclusions derived from downstream transcriptomic analyses? Because cell-type proportions inferred by deconvolution represent model-based estimates rather than direct measurements, uncertainty introduced during deconvolution may propagate into downstream analyses such as differential gene expression, pathway enrichment, and cross-cohort validation. However, the extent to which cell-type correction reshapes disease-associated transcriptional signatures and affects biological interpretation has not been systematically evaluated.

This question is particularly relevant for diseases involving heterogeneous tissues, such as Alzheimer’s disease (AD), a neurodegenerative disorder, where bulk brain transcriptomic studies remain the primary source of large-cohort molecular data. AD pathology is characterized by pronounced alterations in cellular composition, including neuronal loss, reactive gliosis, and microglial activation [10–12]. These disease-associated shifts in cell-type abundance can strongly influence bulk transcriptomic signals, potentially confounding the identification and interpretation of disease-associated genes and pathways across brain regions and cohorts. Although human brain transcriptomic data have been used to benchmark deconvolution algorithms [13] and assess composition-driven effects on differential expression [14], few studies have systematically evaluated how downstream biological interpretations change when cellular composition is explicitly accounted for.

Importantly, while many studies apply deconvolution to correct for cellular composition effects, most analyses implicitly assume that such correction improves biological interpretation. Whether cell-type correction primarily refines disease-associated signals, removes biologically meaningful changes driven by disease-related cellular shifts, or reveals distinct transcriptional programs masked in uncorrected analyses remains unclear.

Understanding this distinction is critical for interpreting transcriptomic studies of heterogeneous tissues such as the brain.

In this study, we therefore focus on systematically assessing the biological impact of cellular deconvolution on downstream transcriptomic analyses. Specifically, we compare disease-associated signals derived from bulk transcriptomic data before and after cell-type correction to determine how deconvolution alters differential gene expression results, pathway-level interpretation, and concordance with genetic risk loci.

To address this question, we analyzed transcriptomic data from two independent Alzheimer’s disease cohorts: the Mount Sinai Brain Bank (MSBB), encompassing four cortical regions— frontal pole (FP), inferior frontal gyrus (IFG), parahippocampal gyrus (PHG), and superior temporal gyrus (STG)—and the ROSMAP cohort, represented by dorsolateral prefrontal cortex (DLPFC) samples. Using dtangle [5] as the primary deconvolution approach, we estimated cell-type proportions and incorporated them into downstream analyses to evaluate how cell-type correction influences differential gene expression patterns, pathway enrichment results, and overlap with AD genome-wide association study (GWAS) loci.

By systematically comparing results obtained before and after cell-type correction across multiple brain regions and cohorts, we establish a framework for distinguishing transcriptional signals driven by changes in cellular composition from those that remain robust after correction. These analyses provide insight into how deconvolution-based correction reshapes disease-associated transcriptomic signatures in heterogeneous tissues and highlight the importance of composition-aware strategies for interpreting bulk transcriptomic data in complex diseases.

## Results

### 1. Overview of our two-pronged analytical framework

An overview of the approach used in this study is described in Fig. 1. Cell-type proportions were first estimated from bulk transcriptomic data using reference-based deconvolution methods such as dtangle and CIBERSORT, and subsequently incorporated as covariates in linear models. Differential expression between control and disease samples was then performed using the edgeR-QLF framework, with and without inclusion of inferred cell-type proportions, while consistently adjusting for technical and biological covariates (e.g., age, RNA integrity number (RIN), post-mortem interval (PMI), and batch). Differentially expressed genes identified under each approach were compared to assess concordance and method-specific signals, followed by downstream functional interpretation and validation.

**Fig. 1.**
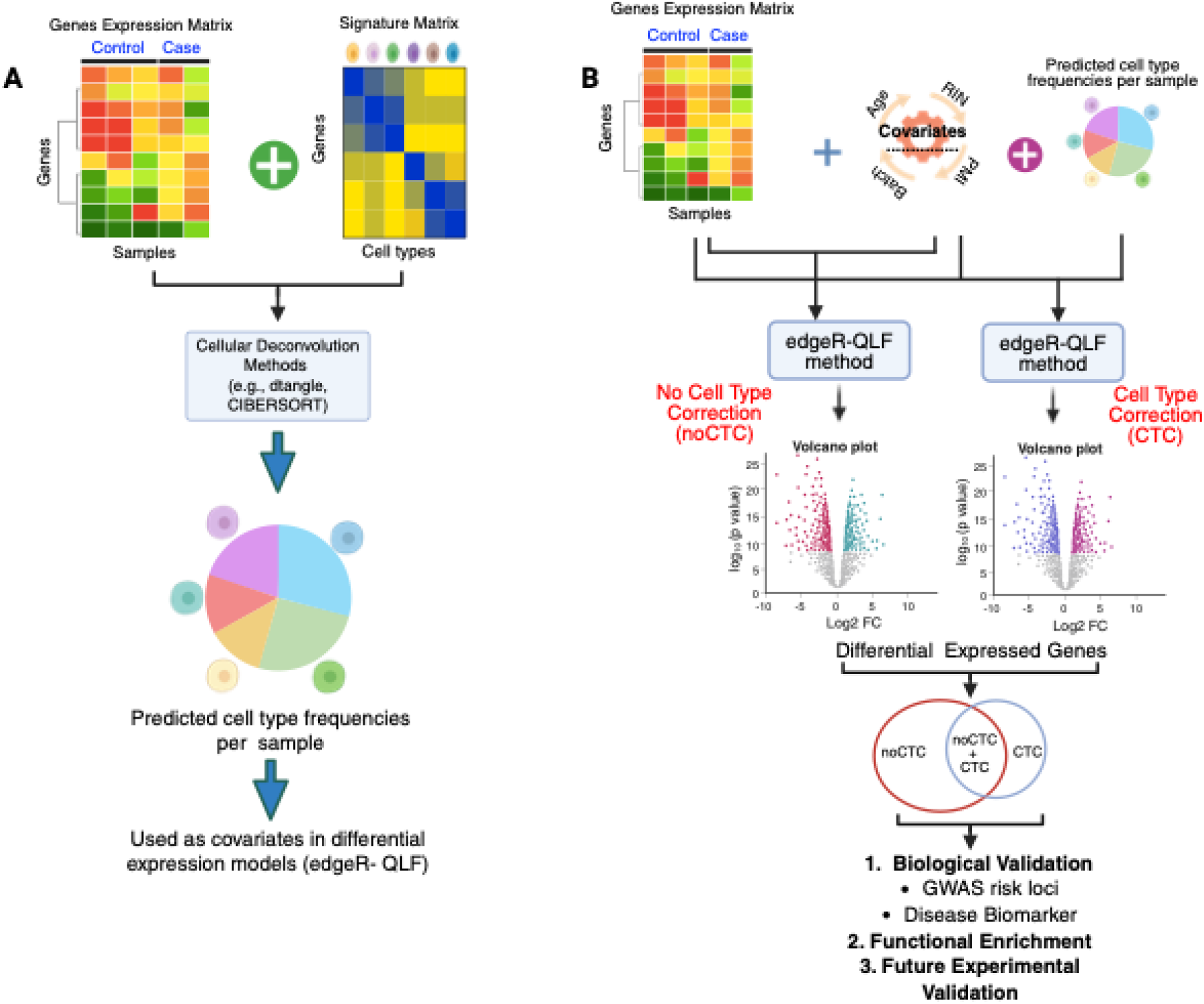
Systematic pipeline to assess the impact of cell-type correction on downstream differential expression analysis. **A** Estimation of cell-type proportions from bulk gene expression data. A bulk gene expression matrix is combined with a reference signature matrix and analysed using reference-based cellular deconvolution methods (e.g., dtangle, CIBERSORT) to infer sample-wise cell-type frequencies. The predicted cell-type proportions are used as covariates in downstream statistical models. **B** Comparative downstream differential expression analysis with and without cell-type correction. Differential expression analysis comparing control (healthy) and case (diseased) samples is performed using the edgeR–QLF method. In both correction modes, technical and biological covariates (e.g., age, RNA integrity number [RIN], post-mortem interval [PMI], and batch) are included. In the cell-type–corrected (CTC) model, predicted cell-type frequencies are additionally incorporated as covariates. Differentially expressed genes (DEGs) identified under each approach are visualized using volcano plots and compared using a Venn diagram to delineate shared and method-specific signals. Resulting gene sets are subjected to downstream biological validation, functional enrichment, and experimental validation.

### 2. Assessing agreement between CIBERSORT and dtangle in terms of their performance

To evaluate the consistency of cell-type correction across deconvolution strategies, we compared CIBERSORT-corrected (CTC^CS^) and dtangle-corrected (CTC^DT^) data by examining both predicted cell-type proportions (Fig. S1A) and log fold change (LogFC) estimates (Fig. S1B) for AD vs control (CTL) differential expression analysis across four brain regions: FP, IFG, PHG, and STG. Scatter plot analyses revealed strong linear concordance between the two methods, with Pearson correlation coefficients ranging from 0.74–0.96 for cell-type proportions and 0.83–0.97 for LogFC estimates across regions (Fig. S1). These results indicate high agreement in both direction and magnitude of differential expression following cell-type correction. The full statistics for Fig. S1B is provided in Table S1.

To further assess method-specific biases and systematic deviations, we employed Bland-Altman analysis [15] (Fig. S1C). The mean differences remained close to zero, indicating no significant systematic bias between the methods. Most genes fell within the 95% limits of agreement, indicating that the methods yield comparable estimates for the majority of DEGs, with only subtle region-specific differences.

Furthermore, dtangle has the advantage of predicting the abundance of a greater number of cell types including rarer cell types such as microglia compared to CIBERSORT, which has been previously shown to estimate the frequencies of abundant cell types such neurons, astrocytes, and oligodendrocytes but not lowly abundant microglia [13,16]. We have used dtangle for all subsequent analyses.

### 3. Validation of cell-type deconvolution and correction of an AD transcriptomic dataset

#### 3.1. Cellular alterations uncovered via deconvolution matches those expected in AD

Cell-type proportion estimates derived using dtangle showed reduced neuronal abundance in AD samples across cortical regions relative to controls, accompanied by corresponding increases in glial populations, including astrocytes, microglia, and oligodendrocytes (Fig. 2A, Fig. S2). These trends are consistent with the established pattern of neuronal vulnerability and reactive gliosis in AD [12]. The magnitude and statistical significance of these compositional changes varied across brain regions (Table S2), indicating region-specific cellular remodeling, with the PHG showing the most pronounced alterations. The observed gradient of neuronal loss and gliosis may reflect the progressive spread of pathology across brain regions.

**Fig. 2.**
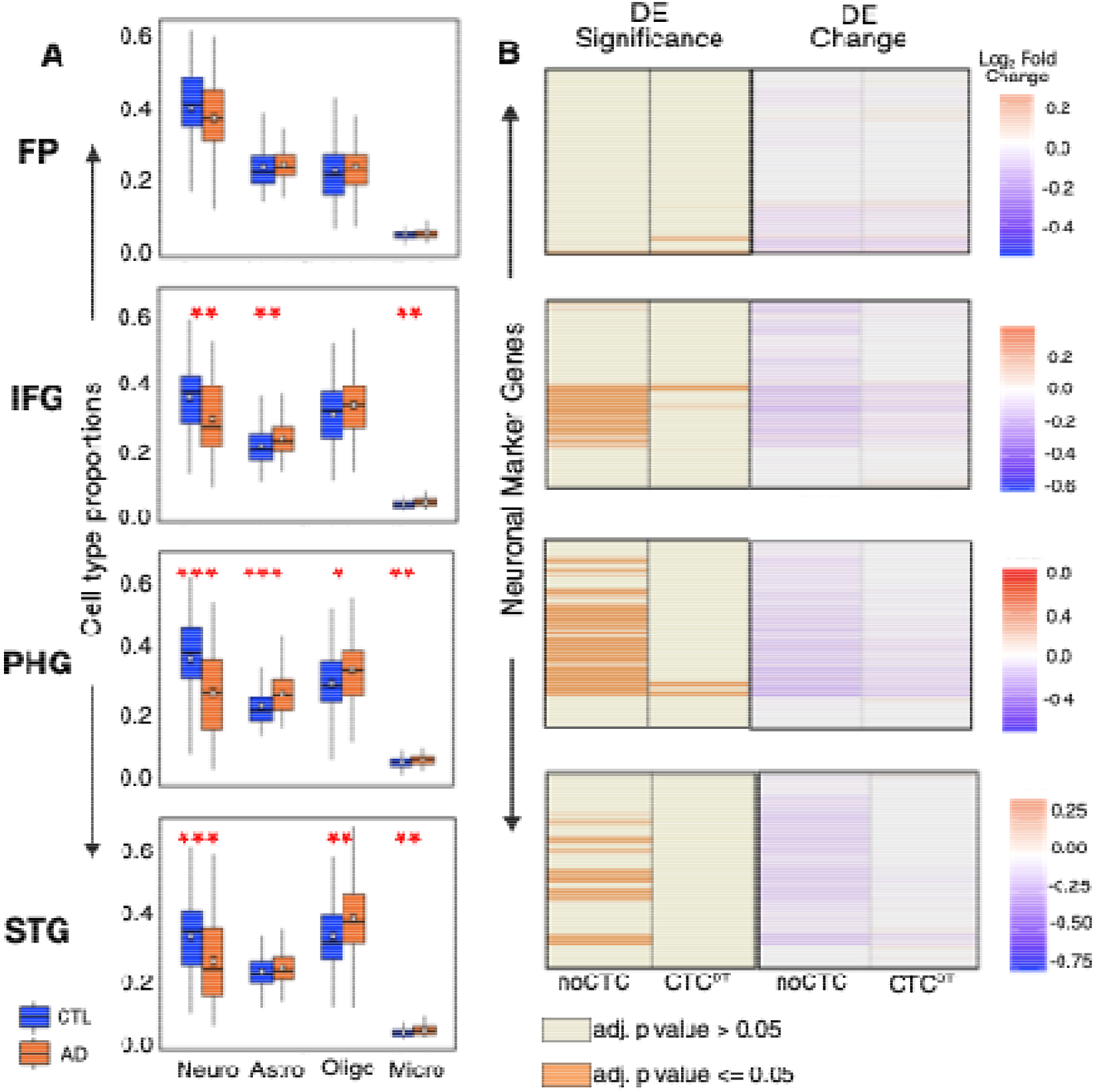
Comparative analysis of cell-type proportions and neuronal marker gene enrichment across brain regions. **A** Boxplots show estimated proportions of major brain cell types across four brain regions, inferred using dtangle. Control (CTL; blue) and Alzheimer’s disease (AD; orange) samples are shown for each region. Statistical differences between AD and CTL samples were assessed independently for each cell type within each brain region using two-sided Wilcoxon rank-sum tests. To account for multiple comparisons, p values were adjusted using the Benjamini–Hochberg false discovery rate (FDR) procedure applied separately within each brain region across the four tested cell types (Neurons, Astrocytes, Oligodendrocytes, and Microglia; *n* = 4 tests per region). Exact p values, adjusted p values (adj. p value) and corresponding effect sizes are reported in Table S2. Statistical significance is indicated using asterisks (adj. *p*□≤□0.05 [*]*, p*□≤□*0.01 [**], and p*□ ≤ □*0.001 [***]),* here and in other figures. Neuro, Astro, Oligo and Micro stands for Neuron, Astrocytes, Oligodendrocytes and Microglia respectively. **B** Two consecutive heatmaps show enrichment patterns of neuron-specific marker genes derived from the MultiBrain signature matrix across brain regions, before (noCTC) and after (CTC^DT^) correction. The left panels (“DE Significance”) show gene-specific adj. p value, with orange indicating adj. p value ≤ 0.05. The right panels (“DE Change”) show the DE effect size as log fold change expressed on a logarithmic scale to the base 2 (log□FC) between AD vs. CTL samples, with purple–red gradients indicating direction and magnitude of change. The DE significance panels show gene-specific adj. p values obtained after Benjamini–Hochberg correction for multiple testing (FDR < 0.05), while the DE change panels display the corresponding log□FC between AD vs. CTL samples. Genes were ordered by adj. p value, and the same ordering was retained across both heatmaps to facilitate direct comparison of significance and effect size patterns before and after correction.

#### 3.2. Overlap with brain marker genes confirms the effectiveness of cell-type correction

To assess whether cell-type-aware correction effectively mitigates composition-driven transcriptional artifacts, we checked the level of significance of the brain cell-type marker genes derived from the MultiBrain signature matrix (covering neurons, astrocytes, microglia, and oligodendrocytes) before correction (noCTC) and after correction (CTC^DT^).

In the noCTC dataset, many cell-type marker genes are significant at FDR 5% particularly in PHG, followed by IFG and STG, across all cell types. This is consistent with the hypothesis that transcriptional changes may be partially driven by shifts in cellular abundance (Fig. 2B, Fig. S3). Even after applying a secondary exclusion filter (retaining only those cell-type marker DEGs not significant in the alternate dataset, p > 0.1; see Table S3), PHG, STG, and IFG preserved the majority of marker genes, while FP retained none. Following correction, there is a clear reduction in the number of significant marker genes in the CTC^DT^ dataset, with STG showing a complete loss of significant marker genes, affirming the removal of composition-driven signals and impact of cell-type correction. In contrast, FP transcriptional signatures remained largely stable following cell-type correction (Fig. 2B, Fig. S3). By comparison, PHG and IFG lost approximately 80–95% of their marker genes following correction (Table 1, Table S3).

We also examined the directionality of fold change for cell-type-specific marker genes (neuronal, astrocytic, microglial, and oligodendrocytic). Consistently across all regions, we observed a downregulation of neuronal markers and an upregulation of glial markers (Fig. 2B, and “marker DE-Change” panels in Fig. S3). As expected, these trends align with dtangle-derived cell-type proportion estimates in AD brains.

These findings, viz., deconvolution outputs matching expected cellular alterations in AD and the pronounced loss of differential expression of marker genes post correction, confirm that CTC^DT^ effectively removes cell-type composition effects. But, noise, as well as interaction between cell-compositional and cell-intrinsic effects, may bias the estimation of cell-type proportions and consequently influence the corrected data. Closer examination at both gene and pathway levels is thus warranted to verify whether cell-type correction refines the DEG landscape toward biologically intrinsic, region-specific transcriptional signatures. To do so, we next assess whether cell-type correction substantially alters the set of DEGs between AD vs CTL, enriched pathways, and related transcriptional signatures associated with AD. By incorporating cell-type relative frequency estimates as covariates in downstream analyses, our evaluation framework allows us to quantify the impact of cellular deconvolution and correction on downstream interpretations, as discussed next.

**Table 1:**
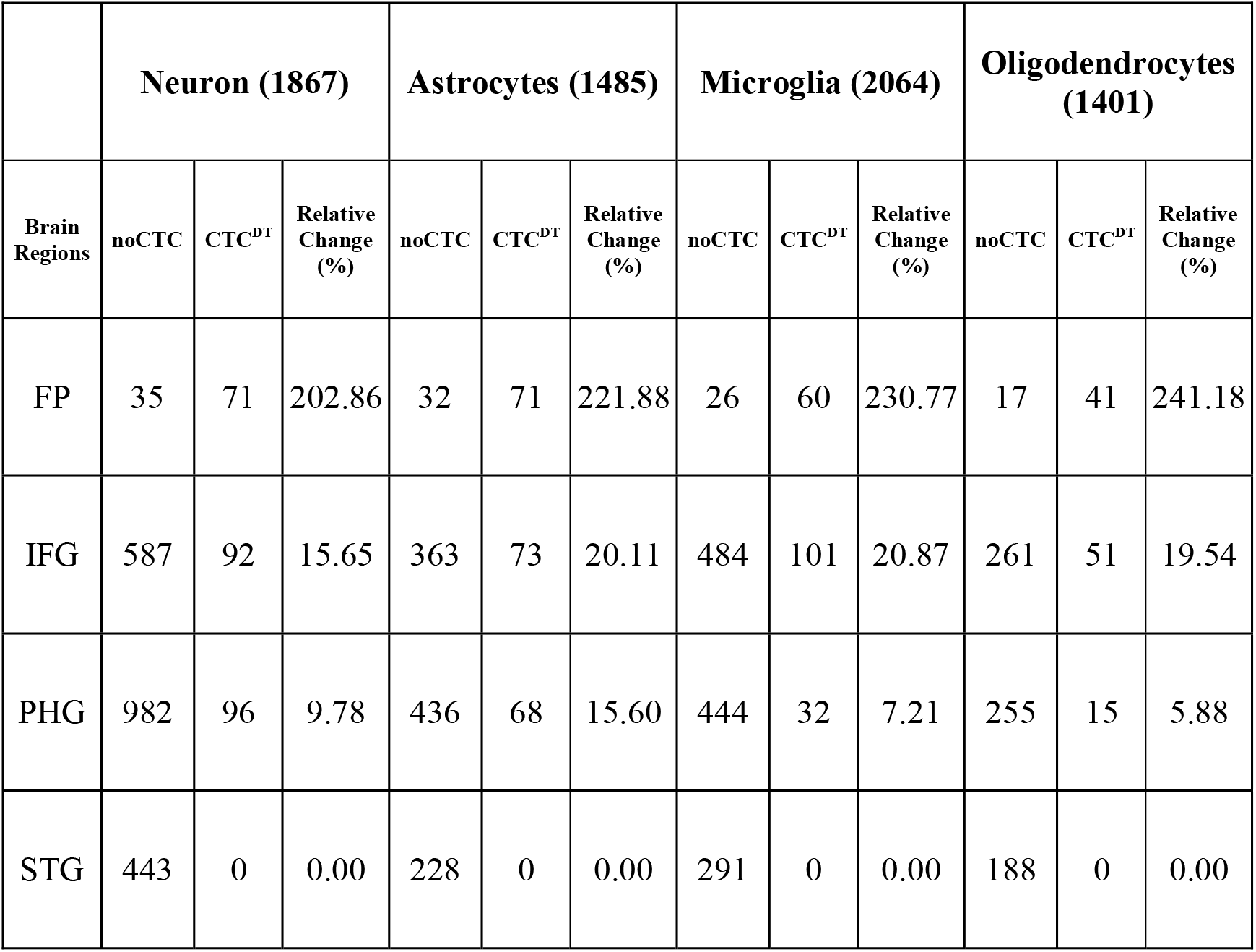
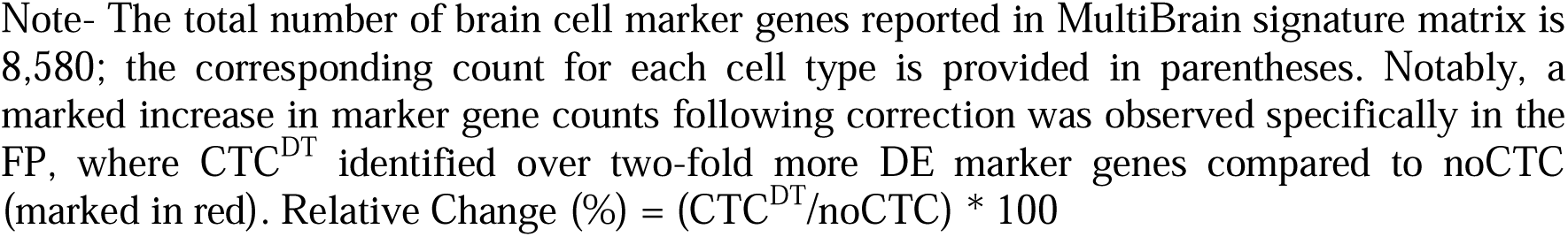
Marker gene DE (status) relative change (%) following CTC^DT^ at FDR 5%. View this in conjunction with the plots shown in Fig. 2B and Fig. S3.

|  | Neuron (1867) |  |  | Astrocytes (1485) |  |  | Microglia (2064) |  |  | Oligodendrocytes (1401) |  |  |
| --- | --- | --- | --- | --- | --- | --- | --- | --- | --- | --- | --- | --- |
| Brain Regions | noCTC | CTC<sup>DT</sup> | Relative Change (%) | noCTC | CTC<sup>DT</sup> | Relative Change (%) | noCTC | CTC<sup>DT</sup> | Relative Change (%) | noCTC | CTC<sup>DT</sup> | Relative Change (%) |
| FP | 35 | 71 | 202.86 | 32 | 71 | 221.88 | 26 | 60 | 230.77 | 17 | 41 | 241.18 |
| IFG | 587 | 92 | 15.65 | 363 | 73 | 20.11 | 484 | 101 | 20.87 | 261 | 51 | 19.54 |
| PHG | 982 | 96 | 9.78 | 436 | 68 | 15.60 | 444 | 32 | 7.21 | 255 | 15 | 5.88 |
| STG | 443 | 0 | 0.00 | 228 | 0 | 0.00 | 291 | 0 | 0.00 | 188 | 0 | 0.00 |

Note- The total number of brain cell marker genes reported in MultiBrain signature matrix is 8,580; the corresponding count for each cell type is provided in parentheses. Notably, a marked increase in marker gene counts following correction was observed specifically in the FP, where CTC<sup>DT</sup> identified over two-fold more DE marker genes compared to noCTC (marked in red). Relative Change (%) = (CTC<sup>DT</sup>/noCTC) \* 100

### 4. Impact of cell-type correction on differential expression analysis

#### 4.1. Exploring transcriptome-wide shifts induced by correction

Cell-type correction leads to transcriptomic shifts of varying magnitude across brain regions, observable even before applying statistical cutoffs. Scatter plots (Fig. 3A) and density plots (Fig. 3B) between noCTC and CTC^DT^ using LogFC values illustrate non-uniform, region-specific remodeling of gene-level significance and fold-change distributions. The full statistics for Fig. 3A is provided in Table S1. These global shifts reflect the extent to which cellular heterogeneity confounds bulk RNA-seq interpretations, demonstrating that neurodegeneration-associated biological signals diffuse to different extents across distinct brain regions.

**Fig. 3.**
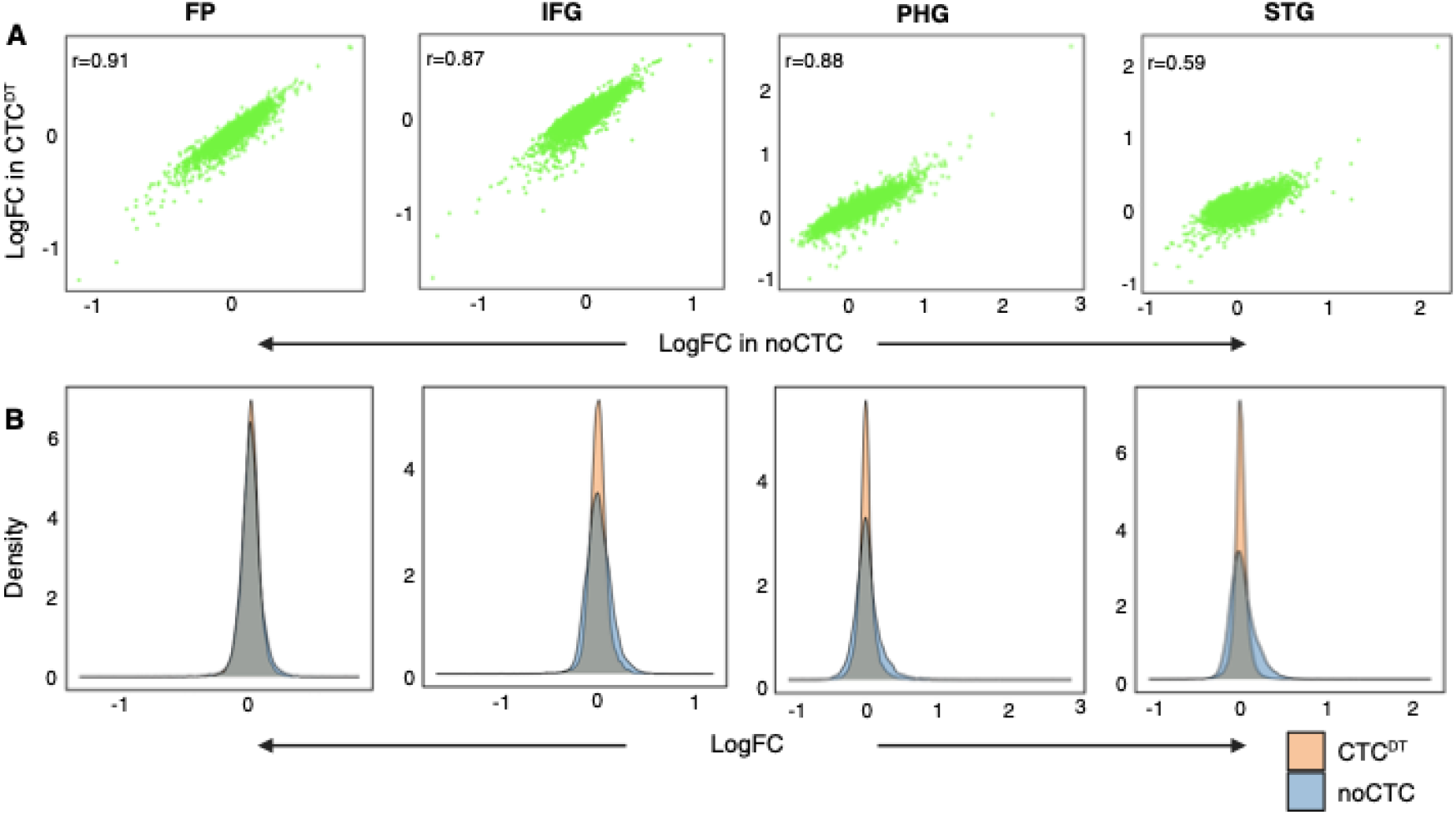
Directional shifts and distributional changes across brain regions post correction. **A** Scatter plots show gene-level LogFC shift across four brain regions, between noCTC (x-axis) and CTC^DT^ (y-axis). Correlation was assessed using both Pearson (r) and Spearman (ρ) coefficients; Pearson r is shown in the plot, and full statistics are provided in Table S1 **B** Each plot reflects the density of LogFC values, highlighting changes in effect size and distribution shape following cell-type-aware correction.

In contrast to the marked reduction in DEG counts observed in IFG and PHG, and the near-complete loss of DEGs in STG following cell-type correction, FP exhibited an increase in DEG detection across all significance thresholds. At FDR ≤ 5%, the number of DEGs increased from 253 in the noCTC dataset to 485 in the CTC^DT^ dataset, corresponding to a 91.7% proportional gain (Table 2). However, the majority of CTC^DT^-associated DEGs substantially overlapped with the noCTC signature (226 shared genes), indicating that correction did not fundamentally alter the underlying transcriptional landscape in FP. Notably, the proportional gain progressively decreased with relaxed significance thresholds, declining from 91.7% at FDR ≤ 5% to 41.1% at FDR ≤ 10% and 26.1% at FDR ≤ 20%. This trend suggests that cell-type correction in FP primarily improves the detection of genes near stringent statistical significance boundaries rather than revealing a substantially distinct disease-associated transcriptional program.

**Table 2:** Impact of deconvolution methods on DEGs identified at FDR 5% varies across brain regions.

| Brain Regions | Sample sizes | DEG count (noCTC) | DEG count (CTC<sup>DT</sup>) | Overlap | ΔDEG | Relative Change (%) |
| --- | --- | --- | --- | --- | --- | --- |
| FP | CTL-106, AD-111 | 253 | 485 | 226 | -232 | 91.7 (Gain) |
| IFG | CTL-87, AD-97 | 3419 | 627 | 576 | 2792 | 81.6 (Loss) |
| PHG | CTL-80, AD-85 | 4439 | 466 | 454 | 3973 | 89.5 (Loss) |
| STG | CTL-85, AD-103 | 2581 | 0 | 0 | 2581 | 100 (Loss) |

Note- $\Delta$ DEG= noCTC-CTC<sup>DT</sup>, Positive $\Delta$ DEG values indicate a reduction in DEG counts after cell-type correction, whereas negative values indicate an increase following correction. Relative change (%) was calculated as: ( $|\Delta$ DEG/noCTC)\*100.
“Gain” indicates increased DEG detection following correction (CTC<sup>DT</sup> > noCTC), whereas “Loss” indicates reduced DEG detection following correction (CTC<sup>DT</sup> < noCTC).
A total of 15,178 genes were analyzed.

In contrast, STG exhibited the most pronounced effect of cell-type correction, with complete loss of significant DEGs at FDR ≤ 5%. Even after relaxing the threshold to FDR ≤ 20%, only seven genes remained significant following correction, indicating that a substantial proportion of the noCTC-associated transcriptional alterations in STG may be driven by changes in cellular composition. Furthermore, although several genes displayed directional reversals between noCTC and CTC^DT^ analyses, none remained significant under the FDR ≤ 5% threshold (Fig. S4).

#### 4.2. Comparison of statistically significant DEGs

Following cell-type correction, DEG counts decline sharply at an FDR threshold of 5%, except in FP—consistent with observations from brain marker gene analyses—reflecting not merely signal refinement but a substantive shift in biological interpretation. CTC^DT^ disentangles disease-associated transcriptional regulation from expression changes driven by cellular composition. A reduction in DEGs is observed in PHG and IFG, while STG shows a complete loss of DEGs, demonstrating that the impact of cell-type deconvolution is region-specific (Table 2).

In FP, IFG, and PHG, most post-correction DEGs were also significant in the uncorrected data, suggesting that a subset of transcriptional signals is resilient to compositional adjustment (see Fig. S5 for the Venn diagram). The identification of genes that remain robust across statistical thresholds and correction approaches highlights them as high-confidence candidates for mechanistic and therapeutic exploration.

We further examined the non-overlapping DEGs identified before and after correction. For each region, we selected genes that were significant in the CTC^DT^ dataset (adj. *p* < 0.05) but not in the uncorrected dataset (*p* > 0.1), with the aim of capturing cell-intrinsic dysregulation in AD (see Methods). This analysis identified 6, 19, and 5 genes in FP, IFG, and PHG, respectively, that were exclusive to CTC^DT^ (see Table S4 for counts of genes that are significant in one condition and demonstrably non-significant in the other). Collectively, this points to potential disease-intrinsic regulatory signals that warrant further functional annotation and pathway-level interrogation, as explored below.

#### 4.3. Impact of cellular heterogeneity on shared transcriptomic signatures

We would like to assess the influence of cellular composition on shared transcriptional signatures across brain regions. To do so, we first identify significant DEGs in each region at FDR ≤ 0.05 using the noCTC (or CTC^DT^) dataset, and then find the significant DEGs shared across the regions to obtain the shared DEGs for the corresponding dataset. 77 DEGs were significant across all four regions (FP, IFG, PHG, and STG) in the noCTC dataset. However, this apparent pan-regional signature was substantially altered following cell-type correction. The significant DEGs shared across regions in the CTC^DT^ dataset were restricted to FP, IFG, and PHG (n = 38 genes), with STG showing a complete loss of significant genes. Notably, the overlap between these 77 and 38 pan-regional DEGs is only 19, indicating that the majority of the initially observed shared transcriptional signals are not preserved after correction (Table S4). Consistent with the shared DEG patterns, pairwise Jaccard similarity between most brain regions decreased substantially following cell-type correction (Fig. S6; Supplementary Text S1).

To further dissect the nature of these shared signals, we performed cell-type enrichment analysis using the MultiBrain signature matrix (Table S5). For this analysis, we first partitioned the union of 77 and 38 pan-regional DEGs detected in noCTC and CTC datasets respectively into three categories: those detected as significant DEGs only in the noCTC dataset (n = 58 genes), those detected both before and after correction (n = 19 genes), and those detected as significant only in the CTC^DT^ dataset (n = 19 genes). Cell-type enrichment analysis revealed distinct biological patterns across these groups. The overlapping genes exhibited significant enrichment for neuronal markers (odds ratio (OR) > 6, adj. p < 0.01), highlighting a robust core of neuron-intrinsic transcriptional changes that are resilient to compositional adjustment. Note that, unless otherwise specified, a statistical significant enrichment discussed here is based on globally adj. p values derived from Benjamini– Hochberg correction across all enrichment tests performed.

The 58 genes detected only in the noCTC dataset showed no significant enrichment, suggesting that they represent weak or heterogeneous signals that are largely mitigated after correction. This pattern appeared to be influenced in part by the FP dataset. Consistent with this interpretation, a subset of noCTC differentially expressed genes (n = 32 genes), present across IFG, PHG, and STG but completely absent in FP, defined by significance in the noCTC dataset (FDR ≤ 0.05) and clear non-significance in the CTC^DT^ dataset (p > 0.1; see Methods), showed significant enrichment for neuronal markers (OR ≈ 3.8, FDR < 0.01). This indicates that a substantial fraction of neuronal signals detected in bulk data arises from changes in cellular composition, such as neuronal loss, rather than intrinsic transcriptional regulation. Consistent with these observations, the broader set of detectable DEGs shared across IFG, PHG, and STG in the noCTC dataset (n = 983 genes) showed strong enrichment for neuronal markers (OR = 2.69, FDR ≈ 7.4e-32), along with comparatively weaker contributions from other cell types, reflecting a dominant but heterogeneous neuronal signal.

Importantly, neuronal enrichment within the noCTC_stringent gene set (n = 32 genes) and the deconvolution-retained overlap set (n = 19 genes) was driven by two distinct subsets of neuronal markers with different regional distributions and gene compositions. The noCTC_stringent group contained neuronal markers such as *DLG3, DNM1L, DZIP3, IPO5, NUDCD3, SERINC1, SLC22A17, SLC9B2, TM2D3, TUSC3, USP33*, which were shared across IFG, PHG, and STG but absent in FP, suggesting region-restricted neuronal signals likely influenced by compositional changes. These genes are consistently significant in PHG, IFG, and STG before correction but consistently lose significance after cell-type correction, while remaining non-significant in FP throughout. In contrast, the CTC-retained group comprised a distinct set of neuronal markers, including *BNIP3, ICA1, KCNN2, MLIP, PPEF1, RIMS1, RIMS2, TASP1, UNC13C*, which remained significantly differentially expressed across all four regions in the noCTC dataset but remained significant (FDR ≤ 0.05) after correction in FP, IFG, and PHG, whereas their significance was largely lost in STG. The distinct composition of neuronal markers between these two groups suggests that they may represent different biological programs, reflecting composition-driven neuronal loss in one case and intrinsic transcriptional dysregulation in the other. This interpretation is consistent with previous studies showing that neuronal modules in Alzheimer’s disease exhibit both differential expression and altered co-expression arising from a combination of neuronal loss and intrinsic regulatory changes [14].

In contrast, genes only detected following correction (n = 19 genes) showed significant enrichment exclusively for astrocyte markers, whereas the broader corrected gene set (n = 38 genes) demonstrated enrichment for both astrocyte and neuronal markers (Table S5), with the neuronal signal primarily contributed by those detected both before and after correction. These observations suggest that cell-type correction not only unmasks astrocyte-associated transcriptional dysregulation but also preserves a subset of neuron-intrinsic transcriptional signals within the shared gene set.

Together, these findings demonstrate that cellular heterogeneity can generate an inflated and composition-driven impression of shared transcriptional activity across brain regions. Cell-type correction effectively disentangles these effects, revealing a smaller, biologically meaningful set of regionally consistent genes and distinguishing between composition-driven neuronal signals and true cell-intrinsic regulatory changes.

### 5. Differential pathway landscapes unveiled by cell-type aware correction-

To deepen our understanding of the biological relevance of transcriptional changes, we performed functional enrichment analysis before and after cell-type correction. We performed Gene Set Enrichment Analysis (GSEA) adopting a cut-off-free approach incorporating all genes and their corresponding LogFC values (see Fig. S7 and Methods for this GSEA workflow).

Heatmap clustering revealed broad regional patterns across both Reactome (Fig. 4A) and Gene Ontology Biological Process (GO_BP) (Fig. 4B). In total, 529 Reactome pathways and 452 GO_BPs were analyzed. PHG and STG consistently clustered together, whereas FP and IFG formed a separate cluster, broadly reflecting segregation between temporal and frontal cortical regions, although minor deviations were observed. Region-specific analyses revealed a consistent reduction in significantly enriched pathways after correction. In PHG and STG, the number of distinct Reactome and GO_BP pathways enriched in the noCTC dataset markedly exceeded those detected in the CTC^DT^ dataset, consistent with the higher number of DEGs identified prior to correction. Exclusive functional heatmaps (Fig. S8) further highlight the functional divergence introduced by deconvolution.

**Fig. 4.**
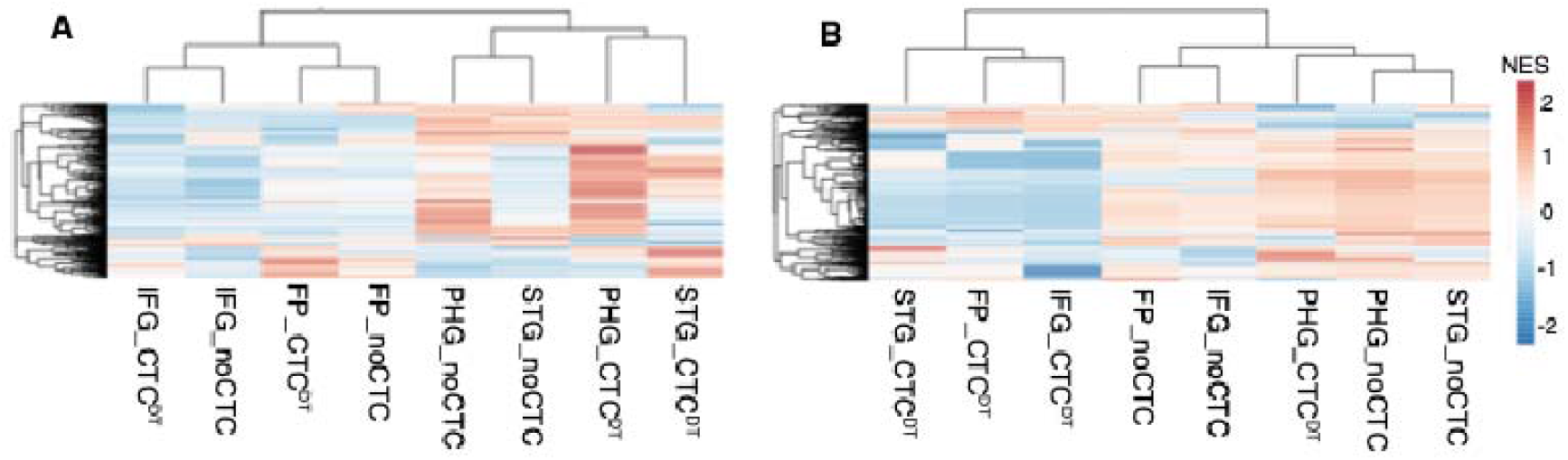
Functional enrichment across brain regions before (noCTC) and after cell-type correction (CTC^DT^). **A** Heatmap of Reactome pathway enrichment across brain regions. Hierarchical clustering separates most noCTC and CTC^DT^ samples, highlighting distinct transcriptional states after cell-type correction. FP and IFG group within one major clade, whereas PHG and STG form another, indicating region-specific functional organization. **B**. Heatmap of Gene Ontology Biological Process (GO_BP) enrichment across brain regions. CTC^DT^ and noCTC samples predominantly form separate clusters; however, corrected PHG (PHG_CTC^DT^) clusters closer to PHG_noCTC, suggesting partial preservation of region-specific biological process signatures following cell-type-aware correction. Colours indicate normalized enrichment scores (NES), ranging from low (blue) to high (red).

In the noCTC dataset, both Reactome and GO_BP demonstrated strong enrichment of neuronal synaptic signaling among AD-downregulated genes, with Reactome pinpointing NMDA receptor activation, long term potentiation, and CREB related cascades, and GO_BP highlighting broader neurotransmission and neuromodulatory processes. In parallel, several classical non-neuronal pathways such as immune signaling, extracellular matrix (ECM) remodeling, coagulation programs appear enriched among AD-upregulated genes, reinforcing the conventional view of AD as a hyper inflammatory and synaptically impaired state. In contrast, following cell-type correction, several immune-related pathways switched direction: interleukin responses, Toll-like receptor cascades, and related innate immune programs were suppressed. These direction changes after proportion adjustment are consistent with the known sensitivity of bulk pathway enrichment to shifting microglial program composition across AD progression [17], and with prior network analyses reporting the emergence of microglial immune-suppression programs (NSupp) in tau-associated neurodegeneration [18].

In addition, CTC^DT^ introduced several DNA repair, replication stress, chromatin regulation categories among AD downregulated genes, pointing to compromised genome maintenance. The amyloid fiber formation pathway remained enriched following cell-type correction, consistent with the well-established role of amyloid pathology in AD. This enrichment involved several genes implicated in APP trafficking and processing, including *SORL1*, *PSENEN*, *FURIN*, *GGA2*, *GGA3*, and *TSPAN14*, [19–21] suggesting that amyloid-related molecular interactions remain perturbed independent of cellular composition changes. Notably, this enrichment was observed specifically in the STG, indicating potential region-specific amyloid-associated network alterations.

Overall, our findings show that many AD-associated signals are driven by altered cellular composition, whereas cell-type-corrected analyses uncover intrinsic deficits in stress-response, innate immune, and synaptic maintenance pathways, emphasizing the importance of accounting for cellular heterogeneity in AD studies.

### 6. Understanding AD-GWAS genes in the context of cell-type correction-

To evaluate the biological relevance of transcriptional changes identified through cell-type– aware correction, we assessed the overlap between DEGs and AD GWAS risk loci. AD risk genes were derived from the Open Targets Platform and filtered using a locus-to-gene (L2G) score threshold (≥ 0.2), yielding 769 GWAS-supported AD genes for downstream analysis (see Methods). These genes were intersected with DEGs identified in each brain region using both noCTC and CTC^DT^ expression data.

As shown in Table S6, the overlap between AD GWAS genes and DEGs (FDR ≤ 5%) recapitulated the global differential expression trends observed earlier. Specifically, the absolute number of overlapping GWAS genes was generally higher in the noCTC data across all regions except the FP, reflecting the substantially larger DEG sets detected prior to correction. The STG showed no overlap following correction, consistent with the absence of significant DEGs in the CTC^DT^ dataset. The PHG exhibited the highest overlap both before (n = 244 genes) and after correction (n = 49 genes).

To determine whether these overlaps exceeded random expectation, we performed hypergeometric enrichment analysis using all tested genes as background. Despite containing fewer DEGs overall, CTC^DT^-associated signatures demonstrated stronger and more specific enrichment for AD GWAS loci compared with noCTC signatures. The strongest enrichment was observed in PHG, where 49 of 466 CTC^DT^-associated DEGs overlapped AD GWAS genes (10.5%; p value < 0.001), whereas the substantially larger noCTC DEG set showed weaker enrichment (244 of 4439 genes; 5.5%; p value = 0.20). Similarly, the DEGs common to noCTC and CTC^DT^ set in PHG also showed significant enrichment (48 of 454 genes; 10.6%; FDR < 0.001). In contrast, enrichment in FP and IFG did not remain significant after multiple-testing correction.

Comparative proportion testing further confirmed that CTC^DT^-associated DEG signatures in PHG contained a significantly higher fraction of AD GWAS genes than the corresponding noCTC signatures (10.5% vs 5.5%; p value = 4.1 × 10), indicating that cell-type correction increases the specificity of disease-relevant transcriptional signals rather than simply reducing DEG counts. A similar but weaker trend was observed in IFG (5.6% vs 3.7%; FDR = 0.047), whereas no significant difference was detected in FP.

A relatively small subset of genes remained significant in both conditions (FP: 15, IFG: 30, PHG: 48, STG: 0), with only five genes—*DUSP6, KCNN2, MSL2, NOX4,* and *RIMS1*— consistently overlapping across FP, IFG, and PHG. Notably, with the exception of *MSL2*, all four genes were also significant AD risk DEGs in the noCTC dataset of STG. Furthermore, although no genes passed the multiple-testing threshold in the CTC^DT^ STG dataset, *DUSP6*, *KCNN2*, and *NOX4* remained significant at a nominal raw p-value cutoff of 5%, suggesting persistence of disease-associated transcriptional signals even after cell-type correction. These genes likely represent robust AD-associated signals whose transcriptional dysregulation cannot be fully attributed to shifts in cell-type composition, suggesting contributions from both cellular proportion changes and cell-intrinsic regulatory mechanisms.

Although these genes are not reported to be known AD biomarkers, their associations with neurodevelopmental, psychiatric, and neurological disorders indicate broader involvement in brain-related pathophysiology and potential relevance to neurodegenerative processes.

Functionally, several of these genes converge on key mechanisms implicated in AD. *RIMS1*, a member of the RAS superfamily, regulates synaptic vesicle exocytosis and neurotransmitter release and has been reported to be downregulated in AD brain tissue (also downregulated in our datasets), potentially contributing to synaptic loss and cognitive decline [22]. Additionally, *RIMS1*-derived circular RNAs (circRNAs) have been shown to exhibit dynamic regulation during AD progression [23]. *KCNN2* encodes a calcium-activated potassium channel that modulates neuronal excitability and synaptic transmission. Collectively, the reduced expression of *RIMS1* and *KCNN2* indicates perturbations in synaptic function and calcium homeostasis, both of which are key hallmarks of Alzheimer’s disease pathology.

*DUSP6* [24], a negative regulator of ERK/MAPK signaling, has been associated with disease severity, with reduced expression correlating with higher clinical dementia rating scores and decreased levels observed in the 5xFAD amyloidopathy mouse model. *NOX4* [25], a key mediator of reactive oxygen species (ROS) production, has been implicated in tau pathology; interestingly, it was consistently downregulated across all regions in our dataset, suggesting potential context-dependent regulation in AD.

In contrast, although no AD risk genes were consistently significant across regions in the CTC^DT^ dataset, a subset retained region-specific significance—*IL6R* in FP, *LAP3* and *ELOVL7* in IFG, and *LILRA5* in PHG. While *IL6R* variants are associated with increased IL-6 pathway activity in Alzheimer’s disease [26], its downregulation in the FP after cell-type correction suggests that elevated bulk signals may reflect cellular composition, particularly immune cell enrichment, rather than cell-intrinsic upregulation. Conversely, its upregulation in the PHG noCTC dataset indicates region-specific inflammatory responses, highlighting spatial heterogeneity in IL-6 signaling. *ELOVL7*, involved in very long-chain fatty acid elongation, points to dysregulation of lipid metabolism, an emerging hallmark of Alzheimer’s disease [27]. Similarly, *LILRA5*, a leukocyte immunoglobulin-like receptor identified as a novel AD risk gene through genetically regulated expression analyses, supports the involvement of immune and microglial pathways in disease pathogenesis [28].

### 7. Cross-cohort validation of differential expression profiles

To assess transcriptional consistency across brain regions and datasets, we compared DEGs between ROSMAP-DLPFC and MSBB-FP. Across both datasets, the expression profiles of 15,131 common genes in DLPFC and FP, formed the basis for cross-region comparisons. Given that DLPFC (Brodmann area 9) and FP (Brodmann area 10) are anatomically adjacent regions within the frontal cortex, we investigated the extent of their shared transcriptional changes.

A comparison using the noCTC data revealed a larger DEG set in DLPFC (3804 genes) compared to FP (253 genes) at FDR ≤ 0.05, with only 86 genes shared between the two regions (Fig. 5A). Although this overlap was statistically significant (p = 9.3 × 10^-4^), the Jaccard Index (JI = 0.02) indicated very low transcriptional concordance across cohorts in the absence of cell-type correction. In contrast, a parallel comparison using CTC^DT^ showed a reduced DEG count in DLPFC (2191) and increased in FP (485), with only 111 genes shared (Fig. 5B). While the statistical significance of overlap improved (p = 3.3 × 10□□), the JI remained low (0.04), indicating minimal intersection between regions at an FDR threshold of 5%. Although cell-type correction improved the statistical significance of overlap and increased the Jaccard Index two-fold, the overall concordance remained low, suggesting limited shared transcriptional programs between regions.

**Fig. 5.**
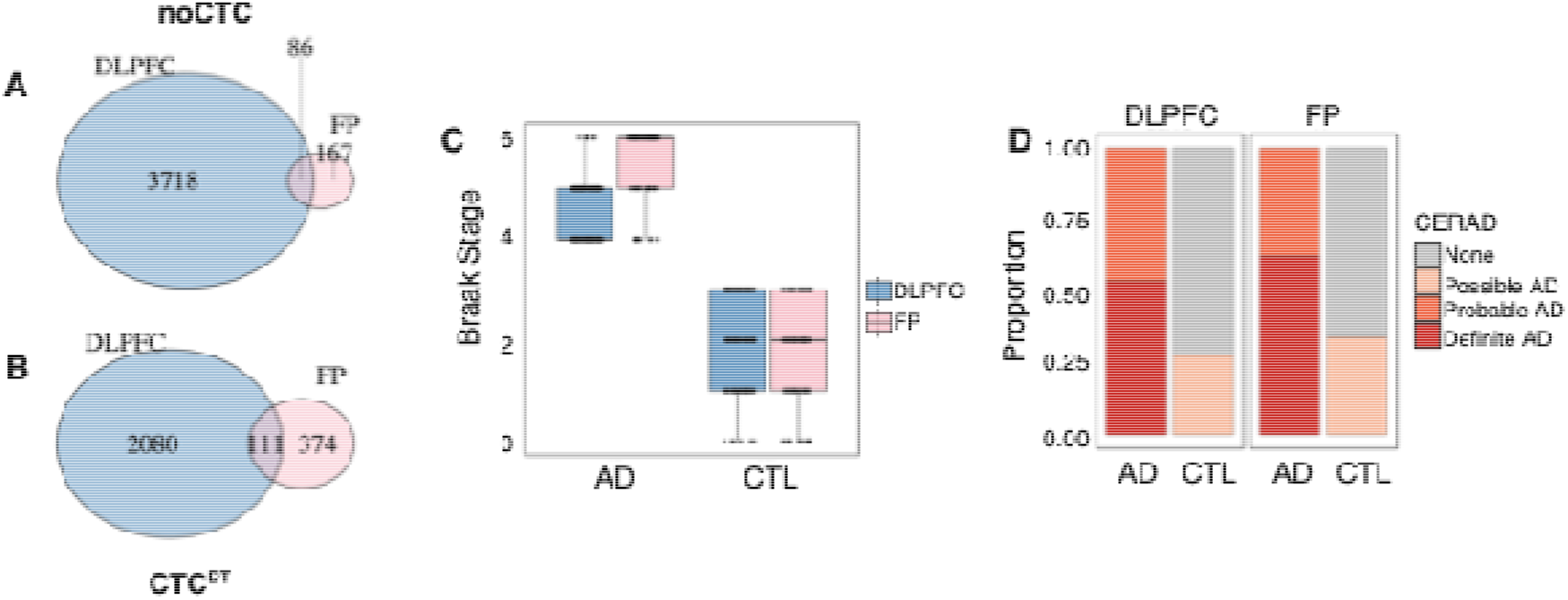
Cross-cohort comparison of DE and disease staging between DLPFC and FP. **A–B** Venn diagrams show the overlap of DEGs between ROSMAP-DLPFC and MSBB-FP under uncorrected (noCTC, A) and cell-type-aware corrected (CTC^DT^, B) conditions. DEG overlap increased following correction, with Jaccard Index (JI) shifting from 0.02 to 0.04; however, overall concordance remained low, indicating limited shared transcriptional signals between regions. **C** Box plots compare Braak staging between FP (MSBB, pink) and DLPFC (ROSMAP, blue) samples. **D** CERAD scoring distributions are represented in this bar plot. AD samples of FP predominantly cluster within Braak stage 6 and CERAD score 2, suggesting later-stage AD pathology relative to DLPFC. These staging differences influence may impact of cell-type correction and underscore the importance of cohort context in transcriptomic analyses, however sample size difference should also be taken under consideration.

Notably, 55 genes were common among the overlapping DEG sets identified in the noCTC (Fig. 5A) and CTC^DT^ (Fig. 5B) comparison, suggesting that they represent a core set of transcriptional alterations conserved across regions and independent of cellular composition. Among these, 4 genes overlap with known AD risk loci, including *NOX4* and *RIMS1*, which were also consistently identified as shared risk-associated genes across MSBB brain regions. This convergence highlights a subset of genes linking genetic susceptibility to conserved, cell-intrinsic transcriptional alterations in AD.

Within-region comparisons further highlighted the differential impact of cell-type correction at FDR ≤ 0.05. In FP, a large proportion of DEGs identified after correction were unique to the corrected analysis (259 genes), with only a small subset detected in the uncorrected condition (27 genes) and 226 genes shared between both approaches (Fig. S5). In contrast, DLPFC showed a marked reduction in DEGs following correction, with a large fraction unique to the uncorrected analysis (2,201 genes), 1,920 genes shared across both conditions, and 474 genes uniquely identified after correction (Fig. S9A). These patterns indicate a stronger influence of cell-type composition on DEG detection in DLPFC compared to FP.

In the DLPFC, of 4,121 DEGs significant without cell-type correction (padjnoCTC ≤ 5%), 530 (∼13%) were lost following correction (padjnoCTC ≤ 5% & pvalCTC^DT^ ≥ 0.1), suggesting a composition-driven contribution; conversely, cell-type correction revealed 2,191 DEGs, including 176 genes uniquely emerging after correction (padjCTC^DT^ ≤ 5% & pvalnoCTC ≥ 0.1) that likely reflect cell-intrinsic transcriptional dysregulation otherwise masked by cellular heterogeneity (Table S7).

To investigate the basis of this low concordance, we examined disease severity distributions across cohorts using Braak staging and CERAD scores (Fig. 5C–D). Notably, FP samples exhibited a higher proportion of advanced pathology, including Braak stage VI and “Definite AD” classifications, compared to DLPFC. FP showed relatively modest and non-significant changes in neuronal and glial proportions, whereas DLPFC demonstrated significant neuronal loss accompanied by corresponding increases in glial populations (Fig. 2A, Fig. S9B). The difference may also be influenced by sample size, as the FP cohort comprised only 217 samples compared with 476 samples in the DLPFC cohort.

Together, these findings indicate that the transcriptional signal in DLPFC is influenced by cell-type composition, particularly neuronal depletion, which inflates DEG detection in uncorrected analyses. In contrast, FP exhibits transcriptional alterations that are less driven by compositional changes and remains stable following cell-type correction. These region-specific differences likely contribute to the limited overlap of DEGs between cohorts, despite both representing AD pathology within the frontal cortex.

## Discussion

In this study, we systematically evaluated the impact of cell-type-aware correction on differential expression, functional enrichment, and cross-cohort consistency across multiple brain regions. Dtangle-based deconvolution uncovered consistent neuronal loss and glial expansion across cortical regions, with PHG exhibiting the most pronounced changes. This confirms the previous reports of PHG being the most vulnerable region experiencing maximum neuronal loss at early stages of diseases [29].

Importantly, cell-type-aware correction markedly reduced the overlap between differentially expressed genes and canonical cell-type markers, in PHG and IFG, and completely eliminated significant DEGs in STG, indicating effective removal of composition-driven signals, with exception in FP. These region-specific shifts in DEGs, together with the emergence of persistently significant gene sets, demonstrate that cell-type-aware deconvolution not only attenuates composition-induced artifacts but also reveals cell-intrinsic regulatory alterations that are obscured in bulk tissue analyses. The behavior of neuronal markers before and after correction provides a clear illustration of this distinction.

Closer examination of these persistently significant gene sets suggests that deconvolution distinguishes two biological components of neuronal pathology: composition-driven neuronal loss and neuron-intrinsic transcriptional dysfunction. One subset of neuronal markers (e.g., *DLG3* and *DNM1L*) lost significance following cell-type correction across PHG, IFG, and STG, consistent with their apparent differential expression being largely driven by neuronal loss and associated changes in cellular composition. In contrast, a second subset of neuronal markers (e.g., *RIMS1*, *RIMS2*, *KCNN2*, and *UNC13C*) remained significantly differentially expressed after correction across PHG, IFG, and FP, suggesting that these genes represent neuron-intrinsic transcriptional dysregulation rather than changes in neuronal abundance. Notably, several of these retained genes are involved in synaptic transmission and neuronal signaling, supporting the view that deconvolution preferentially removes composition-driven neuronal signals while preserving intrinsic molecular alterations in surviving neurons.

Previous benchmarking studies [30] have shown that while deconvolution-based correction improves specificity by reducing compositional confounding, it often does so at the cost of sensitivity, leading to fewer DEGs under stringent thresholds (e.g., FDR ≤ 0.05). This trade-off was consistently observed in our analysis. To mitigate this limitation, we complemented DEG analysis with gene set enrichment analysis (GSEA), which is less dependent on arbitrary significance thresholds. GSEA revealed that the broader enrichment landscape observed in uncorrected data is largely driven by cell-type heterogeneity, capturing signatures such as synaptic dysfunction and immune activation. In contrast, the CTC^DT^-adjusted analysis highlights a more refined transcriptional landscape, characterized by negative enrichment of stress-response and immune regulatory pathways. This shift likely reflects the attenuation of glia-driven inflammatory signals following correction, revealing underlying immune regulatory dysfunction at the cellular level. Taken together, these results underscore a critical distinction. Bulk analyses are strongly influenced by cell-type compositional shifts, inflating apparent immune activation and synaptic downregulation, whereas cell-type–proportion-adjusted analyses instead indicate suppression of stress-response programs and attenuation of innate immune signaling independent of major cell-type proportion differences. This reframes AD pathology as a disorder of cellular redistribution coupled with impaired resilience, where microglial and astrocytic expansion creates the illusion of hyper inflammation, but the true molecular signature is one of suppressed stress responses, weakened innate immune regulation, and enduring synaptic vulnerability. This integrated view emphasizes the importance of cell-type correction in disentangling genuine molecular impairments from compositional shifts, and suggests that therapeutic strategies should focus not only on modulating inflammation but also on restoring intrinsic stress response and synaptic maintenance mechanisms.

Cell-type correction substantially alters the relationship between AD GWAS risk loci and transcriptional changes in the human brain. Although the noCTC data shows a greater absolute overlap between significant DEGs and AD GWAS risk genes, enrichment analyses indicate a stronger and more specific enrichment of AD GWAS risk loci after cell-type correction (CTC^DT^). Notably, only a small subset of genes, including the neuronal markers *KCNN2* and *RIMS1*, remain consistently associated across brain regions following correction, highlighting them as high-confidence candidates that may link AD genetic susceptibility to cell-intrinsic molecular mechanisms.

Cross-cohort comparison between DLPFC and FP revealed unexpectedly low concordance in differential expression despite their anatomical proximity within the frontal cortex. This limited overlap underscores fundamental differences in the underlying drivers of transcriptional change across regions and more likely also study-specific differences such as cohort sample size. In DLPFC, substantial neuronal loss and accompanying glial expansion contribute strongly to differential expression, leading to inflated DEG detection in uncorrected analyses and a marked reduction following cell-type correction. In contrast, FP, despite exhibiting more advanced pathological burden as indicated by Braak staging and CERAD scores, shows relatively stable cellular composition. This decoupling between pathological severity and cellular composition suggests that late-stage disease in certain regions may predominantly reflect cell-intrinsic regulatory alterations rather than overt compositional shifts. Consequently, cell-type correction reduces apparent cross-cohort concordance by removing shared composition-driven signals, while revealing more region-specific regulatory programs. These findings highlight that transcriptional signatures of AD are highly context-dependent, even across adjacent cortical regions, and emphasize that accounting for cellular composition is essential for meaningful cross-cohort comparisons and interpretation of disease-associated gene expression changes.

To evaluate the accuracy and biological fidelity of cell-type correction, we leveraged matched bulk and single-nucleus RNA-seq data [31] from 24 individuals in the ROSMAP cohort. A reference set of seven microglial genes (*CMKLR1*, *VSIG4*, *IFI44L*, *TREM2*, *C1QC*, *FGD3*, *C1orf162*) and seven oligodendrocyte precursor cell genes (*CMTM3, EMILIN3, KIAA0040, PLP1, BTBD17, CRYAB, DDX60*) identified as statistically significant in pseudobulk snRNA-seq analysis served as a benchmark for assessing correction performance [32,33]. This approach is based on the premise that accurate cell-type correction should yield bulk expression profiles that align with the cell-intrinsic patterns identified through pseudobulk snRNA-seq analysis of individual cell types. Consistent with the removal of composition-driven artifacts, when applying actual proportions (CTC^AP^), 2 (*C1QC & VSIG4*) of these 7 microglia genes retain significance (p values < 0.05). When applying dtangle-derived proportions (CTC^DT^), 1 gene (*EMILIN3*) retained significance (p value < 0.05) while the remaining 11 benchmark genes did not reach statistical significance. The differences between dtangle-derived vs. actual proportions and the absence of statistical significance could be attributed to low sample size, and similar analyses could be explored in the future as larger matched single-cell and bulk datasets become available. Future work integrating bulk, single-cell, and spatial transcriptomic approaches will further refine our understanding of AD-associated transcriptional regulation, as these modalities capture complementary population-level, cell-type–specific, and spatial dimensions of gene expression.

Together, our noCTC versus CTC comparison framework presented here demonstrates how cellular deconvolution reshapes the interpretation of AD-associated transcriptional signatures. This approach not only distinguishes compositional from intrinsic disease signals but also prioritizes genes such as *KCNN2* and *RIMS1* that remain robust before and after correction. By systematically separating these components, our workflow provides a principled strategy for refining disease-relevant transcriptional signatures and is readily extendable to other complex diseases characterized by tissue heterogeneity.

## Methods

### Datasets and pre-processing

Metadata and count matrices from different datasets were retrieved from the AD knowledge portal. To categorize samples with pathology as ‘Alzheimer’s’, a Braak (a semiquantitive measure of neurofibrillary tangles) stage of ≥4 and CERAD (a semiquantitive measure of neuritic plaques) label of “DefiniteAD” or “ProbableAD” were considered. Samples were denoted as control (CTL), if they met the criteria of Braak stage ≤ 3, and CERAD labels of “None” or “PossibleAD” [34–36].

**MSBB**-Bulk tissue RNA-seq data from the MSBB (Mount Sinai Brain Bank Study) from the frontal pole and prefrontal cortex (FP and PFC, Brodmann Area 10) region, parahippocampal gyrus (PHG, Brodmann Area 36), inferior frontal gyrus (IFG, Brodmann Area 44), and superior temporal gyrus (STG, Brodmann Area 22) were used in this study. Metadata and count matrices were retrieved from the AD knowledge portal (syn27068754, syn21893059, and syn6101474). Using the criteria of RIN (RNA Integrity Number) scores greater than or equal to 5, we selected 217 samples (211 from FP and 6 from PFC), which comprises 111 AD samples and 106 control samples. For PHG, 165 samples including 85 AD samples and 80 control samples were considered for our study. From IFG, 184 samples including 97 AD samples and 87 control samples were used. Finally, from STG, 188 samples comprising 103 AD samples and 85 control samples were analyzed. 108 samples from all regions belong to the same individuals, mapped based on ‘individualID’.

**ROSMAP**-Bulk tissue RNA-seq data from ROSMAP (Religious Order Study and Memory and Aging Project) were retrieved from the AD knowledge portal (syn8691134, syn21323366, syn3191087, syn21088596 and syn34572333) upon acquiring appropriate approval. We extracted samples from the dorsolateral prefrontal cortex (DLPFC, Brodmann Area 9) region. From the 476 samples, we used samples that had metadata on the age of death, race, RIN scores, PMI (post-mortem interval) and library batch. All selected samples had a RIN score of greater than or equal to 5. Among the 476 samples, there are 295 AD and 181 control samples, defined based on the criteria mentioned earlier, i.e., a Braak stage of ≥4 and CERAD label of “DefiniteAD” or “ProbableAD” as AD and Braak stage ≤ 3, and CERAD labels of “None” or “PossibleAD” as CTL.

Single-nucleus RNA-seq (snRNA-seq) data of 48 ROSMAP samples were obtained from Murphy et al. [37], who had reprocessed data from Mathys et al. [31], with better quality control strategies that included the removal of empty droplets, nuclei with low read counts and doublets. Murphy et al.’s count matrix consists of 50985 cells and 17034 genes. Of these 48 samples, we selected those whose ‘individualID’ could be mapped to any of the 476 ROSMAP samples. 24 such samples were selected, for which all metadata was available, including ‘libraryBatch’. This provided us with paired bulk RNA-seq and snRNA-seq data that were used for DE analysis. The paired samples consist of 9 AD samples and 15 control samples.

We retained only Ensembl gene IDs with a maximum number of counts in all samples, as the count matrices in the bulk dataset consisted of Ensembl gene IDs, while the snRNA-seq count matrix had HGNC gene symbols.

**Pseudobulk snRNA-seq**-We used the ‘pseudobulk’ approach to identify differentially expressed genes for snRNA-seq data, where we aggregate counts to samples, thus accounting for the dependence between an individual’s cells. The analysis was conducted for each distinct cell population, such as microglia, astrocytes, neurons, oligodendrocytes, oligodendrocyte precursor cells, and endothelial cells. Count data from 24 samples is available for Astrocytes, Neurons and Oligodendrocytes. Oligodendrocyte precursor cells, microglia and endothelial cells have data from 22, 22 and 23 samples, respectively. Excitatory and inhibitory neuron cell-type counts are merged and called “neurons”.

## Deconvolution of bulk RNA-seq data

CIBERSORT and dtangle deconvolution algorithms were used to perform cellular deconvolution of bulk RNA-seq data from ROSMAP and MSBB datasets.

Since CIBERSORT and dtangle are reference-based methods, we require a signature matrix to estimate the proportions of all cell types in the samples. For this, we used the multi-brain (MB) signature matrix provided by Sutton et al [13]. This is a composite signature generated by quantile normalising and averaging RPKM (Reads Per Kilobase Million)-level expression of other signatures that were obtained from cortical regions of the brain through various experimental purification protocols for five cell types (neurons, astrocytes, oligodendrocytes, microglia, and endothelial).

CIBERSORT v1.04 was run using the CIBERSORT R package obtained from https://cibersort.stanford.edu. Parameter QN was set to ‘FALSE’ as is recommended in the CIBERSORT portal for RNA-seq data. From the count matrix, rows where the gene counts are less than 50 for all samples put together were removed.

dtangle v2.0.9 R package was used for deconvolution of bulk data. Cell type markers were selected as the top 1% of markers using the ‘find_markers’ function with marker_method = “diff”. Expression data was log (to the base 2) transformed with an offset of 0.5 before computing estimates using the ‘dtangle’ function.

The sample-wise predicted cell-type proportions from CIBERSORT or dtangle can be used as covariates in downstream statistical models/analyses.

## Finding differentially expressed genes

Squair et. al [38], have established that statistical methods edgeR-QLF and edgeR-LRT are least prone to false discoveries in DE analysis. Therefore, we used the edgeR v.4.2.2 R package to fit a quasi-likelihood negative binomial generalized log-linear model to the count data with the glmQLFit function; and conducted differential gene expression analysis between the control (healthy) vs. case (diseased) groups of samples using the glmQLFTest function. Covariates used for ROSMAP data include the age of death, RIN score, PMI, libraryBatch, and gender; in the case of MSBB data, covariates used include the sequencing batch, age of death, RIN score, gender, race, and exonic rate. When conducting differential expression analysis under the cell-type correction mode, predicted cell-type frequencies from CIBERSORT or dtangle are incorporated as additional covariates. The Benjamini-Hochberg (BH) method was used to control the false discovery rate (FDR) using the function p.adjust in R. Statistical significance was assessed at multiple FDR thresholds (5%, 10%, 20%, 30%). We use the gencode.v46 annotations to only assess differentially expressed genes amongst the protein-coding genes.

### A stringent criteria for finding distinct/exclusive DEGs

In several analyses in the manuscript, we require comparison of different sets of DEGs identified in different conditions to identify the shared set of DEGs across these conditions, and importantly also a set of DEGs unique or exclusive to a particular condition. We could obtain these sets by applying simple set operations (i.e., set intersection and set difference) on the appropriate sets of significant genes identified at FDR 5%. However this simple operation may not handle borderline genes correctly as explained next using two conditions A and B. Consider a gene whose adj. p value is 0.001 in condition A and 0.055 in condition B, then this gene may be wrongly classified as an exclusively significant DEG in condition A even though there is some evidence for it to be DEG also in condition B. So we complement our simple set operations based definition of exclusive genes also by a stringent criteria of classifying a gene as exclusive to condition A if its adj. p value in condition A is less than the FDR cutoff of 0.05, and its raw p value in condition B is greater than 0.1.

### Downstream Analyses Functional Enrichment Analysis

To annotate the potential biological functions associated with the DEGs generated from various analyses, we performed GSEA using WebGestalt webserver [39]. Please see. Fig. S7. The complete DEG list for noCTC and CTC^DT^ for all four brain regions (FP, IFG, PHG and STG), along with their corresponding LogFC values, was used for GSEA. Since the ranking statistic was defined as logFC (AD − CTL), the direction of enrichment reflects disease-associated regulation, such that a negative Normalized Enrichment Score (NES) indicates pathways enriched among genes downregulated in AD, whereas a positive NES indicates pathways enriched among genes upregulated in AD. Under the ‘Functional Database Category,’ we selected Gene Ontology Biological Process (GO_BP) and Reactome pathways for enrichment analysis. Initially without applying any cut-off all resulting pathways were considered for testing. In the final step, enrichment results from all eight GSEA runs were combined, retaining only gene sets where at least one of the eight analyses met the significance threshold of FDR ≤ 0.05. Further, only functions exclusively enriched in either noCTC or CTC^DT^ were retained, highlighting transcriptional shifts specific to each analytical framework.

### Selection of AD GWAS Risk Loci

AD risk genes were obtained from the Open Targets Platform (disease ID: MONDO_0004975; API release 25.4.4; accessed December 2025), which integrates multi-source gene–disease association evidence for 12,995 genes. To specifically capture genome-wide association study (GWAS)–derived signals, we used the gwasCredibleSets column corresponding to the Open Targets locus-to-gene (L2G) score. The L2G score is a probabilistic metric (range 0–1) derived from fine-mapped GWAS credible sets and integrated functional genomic evidence, including variant-to-gene distance, eQTL colocalization, and chromatin interaction data to estimate the likelihood that a gene is linked to a GWAS locus.

Of the 12,995 genes in the association table, 1,665 had non-missing L2G scores reflecting fine-mapped GWAS support. Genes lacking L2G evidence were excluded. Among genes with available L2G scores, the median value was 0.164 (mean = 0.234). To enrich for genes with at least moderate L2G support while avoiding overly stringent filtering in a polygenic disease context, we applied a threshold of L2G ≥ 0.2, which lies above the median confidence level of fine-mapped candidates. This filtering yielded a conservative set of 769 AD GWAS-supported risk genes for downstream analyses. Notably, established AD loci such as *APOE*, *CR1*, *TREM2*, *BIN1*, and *CLU* ranked among the highest L2G scores, supporting the biological plausibility of the prioritization strategy.

### Cross-cohort comparison of DE profiles

To assess transcriptional concordance across cohorts and adjacent frontal cortex regions, we compared DEGs identified in ROSMAP-DLPFC and MSBB-FP. Gene identifiers were harmonized using HGNC symbols, and only genes detected in both datasets were retained, yielding 15,131 shared genes that defined the background universe for overlap analyses.

Comparisons were performed separately for noCTC and CTC^DT^ DEG sets at FDR ≤ 5%. For each condition, we quantified the number of DEGs in each dataset and the number of overlapping genes. Statistical significance of overlap was assessed using a hypergeometric test with the 15,131 shared genes as the reference set. Similarity between DEG sets was further quantified using the Jaccard Index (JI), calculated as the ratio of the intersection to the union of the two DEG sets. These analyses were conducted independently for noCTC and CTC^DT^ to evaluate the impact of cell-type correction on cross-cohort concordance.

To contextualize differences in transcriptional overlap, we compared neuropathological measures across cohorts. Braak stage and CERAD categories were obtained for ROSMAP-DLPFC and MSBB-FP samples. Distribution differences were evaluated and visualized to assess variation in disease severity between cohorts. These analyses were performed to determine whether systematic differences in pathological stage could contribute to any difference observed in DEG profiles.

## Declarations

### Availability of data and materials

All code used to perform the analyses and to generate figures in this article can be accessed at https://github.com/BIRDSgroup/Deconvolution-Project/

### Funding

The research presented in this work was supported by Wellcome Trust/DBT grant IA/I/17/2/503323 awarded to M.N.

### Author contributions

S.M. conceived the study and developed its overall framework with inputs from M.N. and M.I. S.M. performed the downstream analyses, data visualization, and biological interpretation of the results, including enrichment, pathway, and GWAS analyses, and prepared the figures and tables with inputs from M.N. and M.I. M.I. performed the cellular deconvolution analyses, contributed to differential gene expression (DGE) analysis, including data preprocessing and comparison of different cellular deconvolution models and methods, and drafted the associated Methods sections with inputs from S.M. and M.N. S.M. wrote the manuscript with contributions and revisions from M.I. and M.N. M.N. provided overall guidance and supervision for the computational and biological aspects of the study. S.M. and M.I. contributed equally to this work.

## Acknowledgments

The authors would like to thank Ishaan Gupta from Indian Institute of Technology Delhi for his valuable comments and suggestions during S.M.’s presentation at IIT Delhi. The authors also gratefully acknowledge the members of the BIRDS (Bioinformatics and Integrative Data Science) lab for their insightful discussions and constructive feedback during group meetings, which helped improve this work. The authors acknowledge the use of Grammarly and ChatGPT solely for language editing and copy-editing assistance.

## Ethics approval and consent to participate

Not applicable

## Consent for publication

Not applicable

## Competing interests

The authors declare no competing interests.

## Notes

### Competing Interest Statement

The authors have declared no competing interest.

