## Supplementary Documents for "Systematic assessment of the biological impact of cellular deconvolution on downstream analyses of disease transcriptomes"

### Table of Contents:

#### Supplementary Text

**Text S1**: Cell-type correction affects cross-region transcriptomic similarity

#### Supplementary Tables

- **Table S1**: Pairwise correlation of log fold-change (logFC) estimates across deconvolution methods ($CTC^{CS}$ vs $CTC^{DT}$) and $CTC^{DT}$ vs noCTC for each brain region.
- **Table S2**: Cell-type proportion differences between AD vs. CTL samples across brain regions.
- **Table S3**: Overlap with brain marker genes at stringent condition confirms the effectiveness of CTC
- **Table S4**: Composition-driven and cell-intrinsic DEG partitioning between noCTC and $CTC^{DT}$ analyses across brain regions
- **Table S5**: Cell-type enrichment of gene sets defined by overlap and stringency across noCTC and $CTC^{DT}$ datasets.
- **Table S6**: Overlap of AD GWAS risk loci with DEGs before and after cell-type correction
- **Table S7**: Composition-driven and cell-intrinsic DEG partitioning in the DLPFC from the ROSMAP replication cohort

#### Supplementary Figures

- **Fig. S1**: Assessing agreement between CIBERSORT and dtangle cellular deconvolution methods across brain regions.
- **Fig. S2**: Comparative analysis of cell-type proportions across four brain regions using matched samples from 108 individuals
- **Fig. S3**: Marker gene enrichment across brain regions.
- **Fig. S4**: Scatter plots show gene-level LogFC across four brain regions, comparing noCTC vs. $CTC^{DT}$.
- **Fig. S5**: Venn diagrams illustrate the overlap of DEGs between noCTC and $CTC^{DT}$.
- **Fig. S6**: Inter-regional transcriptomic relationships using Jaccard similarity indices.
- **Fig. S7**: Computational workflow for functional enrichment of transcriptional changes in Alzheimer's disease.
- **Fig. S8**: Exclusive functional enrichment across brain regions before and after cell-type correction.
- **Fig. S9**: Cell-type proportion estimates and DEG overlap in ROSMAP-DLPFC.

### Supplementary Text

#### Text S1: Cell-type correction affects cross-region transcriptomic similarity

To dissect inter-regional transcriptomic relationships, we computed Jaccard similarity indices to capture patterns of convergence or divergence in DEG profiles at FDR 5% (Fig. S6). The split heatmap illustrates distinct patterns of inter-regional similarity across the two conditions. In the noCTC dataset (lower triangle), moderate similarity was observed between PHG and STG (Jaccard = 0.32), IFG and STG (Jaccard = 0.28), and IFG and PHG (Jaccard = 0.25), supporting the presence of a broadly shared transcriptional signal. In contrast, FP exhibited consistently low similarity with other regions (Jaccard ≤ 0.05), suggesting a distinct transcriptional profile prior to correction.

Following cell-type correction (upper triangle), cross-region similarity decreased markedly across nearly all region pairs. The overlap between IFG and PHG dropped substantially (from 0.25 to 0.07), and PHG–STG and IFG–STG similarities were reduced to zero, consistent with the complete loss of DEGs in STG after correction. These changes indicate that much of the apparent inter-regional concordance in the noCTC dataset is driven by shared cellular composition rather than intrinsic transcriptional regulation.

Notably, FP–IFG and FP-PHG the only two pairs showing an increase in similarity following correction (from 0.05 to 0.16, 0.03 to 0.08 respectively), suggesting that cell-type correction can enhance the detection of shared, cell-intrinsic transcriptional signals that are otherwise masked by compositional effects. This observation is consistent with the increase in significant DEGs in the $CTC^{DT}$ dataset in FP. The increased similarity between FP and IFG post-correction may reflect shared frontal cortical biology that is obscured prior to correction. To ensure that the observed pattern in cross-regional similarity was not an artifact of stringent DEG filtering, we relaxed the significance threshold from FDR 5% to FDR 20%. Despite the increased sensitivity and higher DEG counts, the overall pattern persisted.

Overall, these results highlight strong region-specificity of DEG signatures following correction, with FP appearing more resilient to correction effects, whereas STG exhibits the highest sensitivity. Collectively, these findings demonstrate that cell-type correction substantially reshapes the landscape of cross-region transcriptomic similarity. While bulk data suggest widespread inter-regional concordance, much of this signal is attributable to cellular heterogeneity. Correction reduces spurious overlap and reveals a more restricted set of biologically meaningful, regionally consistent transcriptional signatures.

### Supplementary Tables

**Table S1: Pairwise correlation of log fold-change (logFC) estimates across deconvolution methods ($CTC^{CS}$ vs $CTC^{DT}$) and $CTC^{DT}$ vs noCTC for each brain region.** Pearson (r) and Spearman (ρ) correlation coefficients quantify agreement in effect size and rank, respectively. Direction agreement represents the proportion of genes with consistent direction of change between comparisons. All correlations were statistically significant ($p < 2.2e\text{-}16$). Notably, the comparatively low correlation observed for the $CTC^{DT}$ vs. noCTC comparison in the STG suggests a pronounced impact of cellular deconvolution on differential expression estimates in this region. The scatter plots corresponding to this table are presented in Fig S1B ($CTC^{CS}$ vs $CTC^{DT}$) and Fig. 3A ($CTC^{DT}$ vs noCTC).

| Brain Regions | LogFC Comparison | Spearman (rho) | Pearson (r) | Direction agreement |
|---|---|---|---|---|
| **FP** | $CTC^{CS}$ vs $CTC^{DT}$ | 0.90 | 0.87 | 0.89 |
| | $CTC^{DT}$ vs noCTC | 0.91 | 0.91 | 0.89 |
| **IFG** | $CTC^{CS}$ vs $CTC^{DT}$ | 0.97 | 0.97 | 0.93 |
| | $CTC^{DT}$ vs noCTC | 0.86 | 0.87 | 0.85 |
| **PHG** | $CTC^{CS}$ vs $CTC^{DT}$ | 0.95 | 0.96 | 0.91 |
| | $CTC^{DT}$ vs noCTC | 0.86 | 0.88 | 0.84 |
| **STG** | $CTC^{CS}$ vs $CTC^{DT}$ | 0.84 | 0.83 | 0.85 |
| | $CTC^{DT}$ vs noCTC | 0.50 | 0.59 | 0.68 |

**Table S2: Cell-type proportion differences between AD vs. CTL samples across brain regions.** Statistical comparison of estimated cell-type proportions between AD vs. CTL samples across brain regions. Differences were assessed using two-sided Wilcoxon rank-sum tests with Benjamini–Hochberg correction. Effect sizes are reported as rank-biserial correlation, with negative values indicating reduced and positive values indicating increased proportions in AD. Statistical significance is indicated using asterisks (adjusted $p \leq 0.05$ [*], $p \leq 0.01$ *[**]*, $p \leq 0.001$ *[***] and* $p > 0.05$ *[ns]).* The corresponding figure is represented in Figure 2A.

| Brain Regions | Cell Type | P Value | Adjusted P Value | Effect Size | Significance |
|---|---|---|---|---|---|
| FP | Neurons | 0.02 | 0.08 | -0.18 | ns |
| | Astrocytes | 0.07 | 0.09 | 0.14 | ns |
| | Oligodendrocytes | 0.12 | 0.12 | 0.12 | ns |
| | Microglia | 0.07 | 0.09 | 0.14 | ns |
| IFG | Neurons | 0.0002 | 0.001 | -0.31 | ** |
| | Astrocytes | 0.001 | 0.002 | 0.27 | ** |
| | Oligodendrocytes | 0.13 | 0.13 | 0.13 | ns |
| | Microglia | 0.001 | 0.002 | 0.27 | ** |
| PHG | Neurons | 1.9E-6 | 7.6E-6 | -0.43 | *** |
| | Astrocytes | 2.7E-5 | 5.4E-5 | 0.38 | *** |
| | Oligodendrocytes | 0.03 | 0.03 | 0.19 | * |
| | Microglia | 0.001 | 0.002 | 0.28 | ** |
| STG | Neurons | 5.6E-5 | 0.0002 | -0.34 | *** |
| | Astrocytes | 0.18 | 0.18 | 0.11 | ns |
| | Oligodendrocytes | 7.3E-4 | 0.001 | 0.29 | ** |
| | Microglia | 0.003 | 0.004 | 0.25 | ** |

**Table S3: Overlap with brain marker genes at stringent condition confirms the effectiveness of CTC**

| | Neuron (1867) | | | | Astrocytes (1485) | | | |
|---|---|---|---|---|---|---|---|---|
| | noCTC | | $CTC^{DT}$ | | noCTC | | $CTC^{DT}$ | |
| Brain Regions | $padj_{noCTC} \le 5\%$ | $padj_{noCTC} \le 5\%$ & $pval_{CTC}^{DT} \ge 0.1$ | $padj_{CTC}^{DT} \le 5\%$ | $padj_{CTC}^{DT} \le 5\%$ & $pval_{noCTC} \ge 0.1$ | $padj_{noCTC} \le 5\%$ | $padj_{noCTC} \le 5\%$ & $pval_{CTC}^{DT} \ge 0.1$ | $padj_{CTC}^{DT} \le 5\%$ | $padj_{CTC}^{DT} \le 5\%$ & $pval_{noCTC} \ge 0.1$ |
| FP | 35 | 0 | 71 | 0 | 32 | 0 | 71 | 0 |
| IFG | 587 | 76 | 92 | 2 | 363 | 42 | 73 | 2 |
| PHG | 982 | 365 | 96 | 0 | 436 | 160 | 68 | 1 |
| STG | 443 | 335 | 0 | 0 | 228 | 142 | 0 | 0 |

| | Microglia (2064) | | | | Oligodendrocytes (1401) | | | |
|---|---|---|---|---|---|---|---|---|
| | noCTC | | $CTC^{DT}$ | | noCTC | | $CTC^{DT}$ | |
| Brain Regions | $padj_{noCTC} \le 5\%$ | $padj_{noCTC} \le 5\%$ & $pval_{CTC}^{DT} \ge 0.1$ | $padj_{CTC}^{DT} \le 5\%$ | $padj_{CTC}^{DT} \le 5\%$ & $pval_{noCTC} \ge 0.1$ | $padj_{noCTC} \le 5\%$ | $padj_{noCTC} \le 5\%$ & $pval_{CTC}^{DT} \ge 0.1$ | $padj_{CTC}^{DT} \le 5\%$ | $padj_{CTC}^{DT} \le 5\%$ & $pval_{noCTC} \ge 0.1$ |
| FP | 26 | 0 | 60 | 0 | 17 | 0 | 41 | 1 |
| IFG | 484 | 35 | 101 | 2 | 261 | 18 | 51 | 0 |
| PHG | 444 | 164 | 32 | 1 | 255 | 99 | 15 | 1 |
| STG | 291 | 175 | 0 | 0 | 188 | 128 | 0 | 0 |

*Note: Within each analytical state (such as uncorrected or noCTC, “$padj_{noCTC}$” refers to genes identified as differentially expressed at an FDR ≤ 5% using noCTC data. “$pval_{CTC}^{DT}$” indicates that these same genes do not reach nominal significance (p value ≥ 0.1) in the alternate state (corrected or $CTC^{DT}$). The same logic applies reciprocally to $CTC^{DT}$-derived DEGs evaluated against noCTC. See Methods for details about stringent DE analysis.
The total number of brain cell marker genes reported in MultiBrain signature matrix is 8,580; the corresponding counts for each cell type are provided in parentheses.

**Table S4: Composition-driven and cell-intrinsic DEG partitioning between noCTC and $CTC^{DT}$ analyses across brain regions**

| Brain Regions | noCTC | | $CTC^{DT}$ | |
|---|---|---|---|---|
| | $padj_{noCTC} \leq$ 5% | $padj_{noCTC} \leq$ 5% & $pval_{CTC}^{DT} \geq$ 0.1 | $padj_{CTC}^{DT} \leq$ 5% | $padj_{CTC}^{DT} \leq$ 5% & $pval_{noCTC} \geq$ 0.1 |
| **FP** | 253 | 0 | 485 | 6 |
| **IFG** | 3419 | 298 | 627 | 19 |
| **PHG** | 4439 | 1576 | 466 | 5 |
| **STG** | 2581 | 1682 | 0 | 0 |
| **Common** | 77 | 32 | 38 | 0 |

*Note: Within each state (such as uncorrected or noCTC, "$padj_{noCTC}$" refers to genes identified as differentially expressed at an FDR ≤ 5% for noTC data. "$pval_{CTC}^{DT}$" indicates that these same genes do not reach nominal significance (p value ≥ 0.1) in the alternate state (corrected or $CTC^{DT}$). The same logic applies reciprocally to $CTC^{DT}$-derived DEGs evaluated against noCTC. See Methods for details about stringent DE analysis.

**Table S5: Cell-type enrichment of gene sets defined by overlap and stringency across noCTC and $CTC^{DT}$ datasets.**

| Query Gene Set | Query Size | Regions | Key Enrichment | OR (no. of intersecting genes between query and MultiBrain marker set) | Global adjp | Biological Interpretation |
|---|---|---|---|---|---|---|
| noCTC_4 | 77 | FP ∩ IFG ∩ PHG ∩ STG | none (weak neuronal) | ~1.9 (16) | ns | diluted pan-regional signal |
| noCTC_3 | 983 | IFG ∩ PHG ∩ STG | Neuron | 2.69 (252) | <1e-30 | broad neuronal-dominated signal |
| noCTC-only | 58 | noCTC(FP ∩ IFG ∩ PHG ∩ STG) not in $CTC^{DT}$_3 | none | — | ns | weak/heterogeneous, removed |
| noCTC_stringent | 32 | IFG ∩ PHG ∩ STG not in $CTC^{DT}$_3 | Neuron | ~3.8 (11) | <0.01 | composition-driven neuronal |
| Overlap | 19 | $CTC^{DT}$_3 ∩ noCTC_4 | Neuron | >6 (9) | <0.001 | intrinsic neuronal core |
| $CTC^{DT}$_3 | 38 | FP ∩ IFG ∩ PHG | Astrocyte + Neuron | ~3 (10 + 11) | <0.01 | refined biology |
| $CTC^{DT}$-only | 19 | $CTC^{DT}$_3 − noCTC_4 | Astrocyte | >5 (7) | <0.01 | astrocyte-specific signal |

Note: *noCTC_4*: genes significant (FDR ≤ 0.05) across FP, IFG, PHG, STG; *noCTC_3*: across IFG, PHG, STG; *noCTC-only*: uncorrected-specific genes; *noCTC_stringent*: significant in noCTC but non-significant after correction (p > 0.1); *Overlap*: common to noCTC_4 and $CTC^{DT}$_3; *$CTC^{DT}$_3:*

significant after correction (FP, IFG, PHG); *CTC$^{DT}$-only*: correction-specific genes. Enrichment based on MultiBrain markers; Odds Ratio (OR) and False Discovery Rate (FDR) shown. Gene sets were defined based on differential expression (FDR ≤ 0.05) and cross-condition criteria (p ≥ 0.1), as described in Table S4 and Methods.

Further, although STG failed to retain significant shared DEGs after correction, 17 of the 19 overlap genes remained nominally significant in STG (raw $p < 0.1$), compared with 12 of 19 CTC$^{DT}$-only genes, indicating partial preservation of the neuron-intrinsic shared signature despite reduced statistical power after deconvolution.

**Table S6: Overlap of AD GWAS risk loci with DEGs before and after cell-type correction**

| Brain Regions | noCTC | CTC$^{DT}$ | Common |
|---|---|---|---|
| FP | 17 | 33 | 15 |
| IFG | 128 | 35 | 30 |
| PHG | 244 | 49 | 48 |
| STG | 140 | 0 | 0 |

| FP_IFG_PHG common genes | Gene Full Name |
|---|---|
| *DUSP6* | dual specificity phosphatase 6 |
| *KCNN2* | potassium calcium-activated channel subfamily N member 2 |
| *MSL2* | MSL complex subunit 2 |
| *NOX4* | NADPH oxidase 4 |
| *RIMS1* | regulating synaptic membrane exocytosis 1 |

The table summarizes the number of AD GWAS-supported risk genes (n= 769), derived from Open Targets Platform, that overlap with DEGs across four brain regions—FP, IFG, PHG and STG. The *Common* column indicates genes that are significantly differentially expressed (FDR ≤ 0.05) in both noCTC as well as CTC$^{DT}$ analyses, representing AD GWAS loci whose transcriptional dysregulation persists after accounting for cell-type heterogeneity. Upon further analysing these common genes, we observed that 5 genes are common for FP, IFG and PHG. Out of these 5 genes except *MSL2*, all 4 genes are significant in the noCTC dataset of STG.

**Table S7: Composition-driven and cell-intrinsic DEG partitioning in the DLPFC from the ROSMAP replication cohort**

| **Brain Region** | **noCTC** | | **$CTC^{DT}$** | |
|---|---|---|---|---|
| | $padj_{noCTC} \leq 5\%$ | $padj_{noCTC} \leq 5\%$ & $pval_{CTC}^{DT} \geq 0.1$ | $padj_{CTC}^{DT} \leq 5\%$ | $padj_{CTC}^{DT} \leq 5\%$ & $pval_{noCTC} \geq 0.1$ |
| **DLPFC** | 4121 | 530 | 2191 | 176 |

Note: Same notation as Table S3 and S4 is followed.

<u>**Supplementary Figures**</u>

**Supplementary Figure 1**

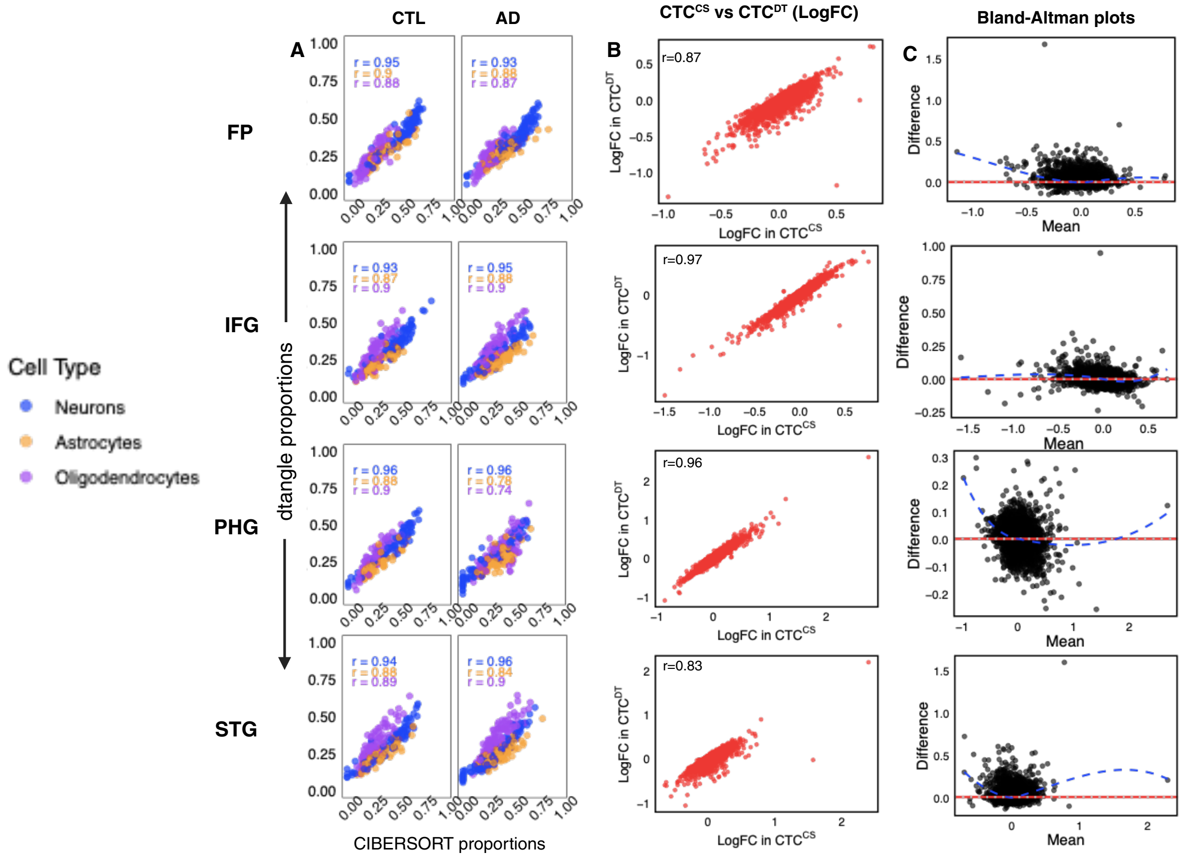


**Fig. S1: Assessing agreement between CIBERSORT and dtangle cellular deconvolution methods across brain regions.** (A) Scatter plots show the relationship between CIBERSORT-estimated (x-axis) and dtangle-estimated (y-axis) cell-type proportions for neurons, astrocytes, and oligodendrocytes between AD vs CTL derived from gene expression data of four brain regions. Estimates are shown separately for control (CTL; left side and Alzheimer's disease (AD; right side) samples. (B) Comparison between gene-wise log fold-change (logFC) estimates based on CIBERSORT ($CTC^{CS}$) and dtangle ($CTC^{DT}$) methods yields highly correlated DE results (Table S1). (C) Bland-Altman plots assessing agreement between the two methods further corroborates the similarity between two methods. Difference stands for "Difference between LogFC of $CTC^{CS}$ and $CTC^{DT}$" and Mean stands for "Mean of LogFC of $CTC^{CS}$ and $CTC^{DT}$". Most genes fell within the 95% limits of agreement, indicating that the methods yield comparable estimates for the majority of DEGs. However, residual plots revealed region-specific deviations. In FP, residuals were randomly scattered around zero, consistent with homoscedasticity and a well-calibrated linear fit. In IFG and PHG, residuals exhibited wave-like and funnel-shaped patterns, respectively, suggesting non-linear effects and heteroscedasticity in subsets of genes, possibly due to differential sensitivity of the methods to transcriptomic complexity or cell-type composition. Interestingly, in STG, the residual trend line dipped on the right-hand side, indicating a tendency for $CTC^{DT}$ to produce slightly lower LogFC values for highly expressed genes compared to $CTC^{CS}$. This asymmetric bias may reflect region-specific transcriptional architecture or differential performance of the methods in handling high-abundance transcripts.

**Supplementary Figure 2**

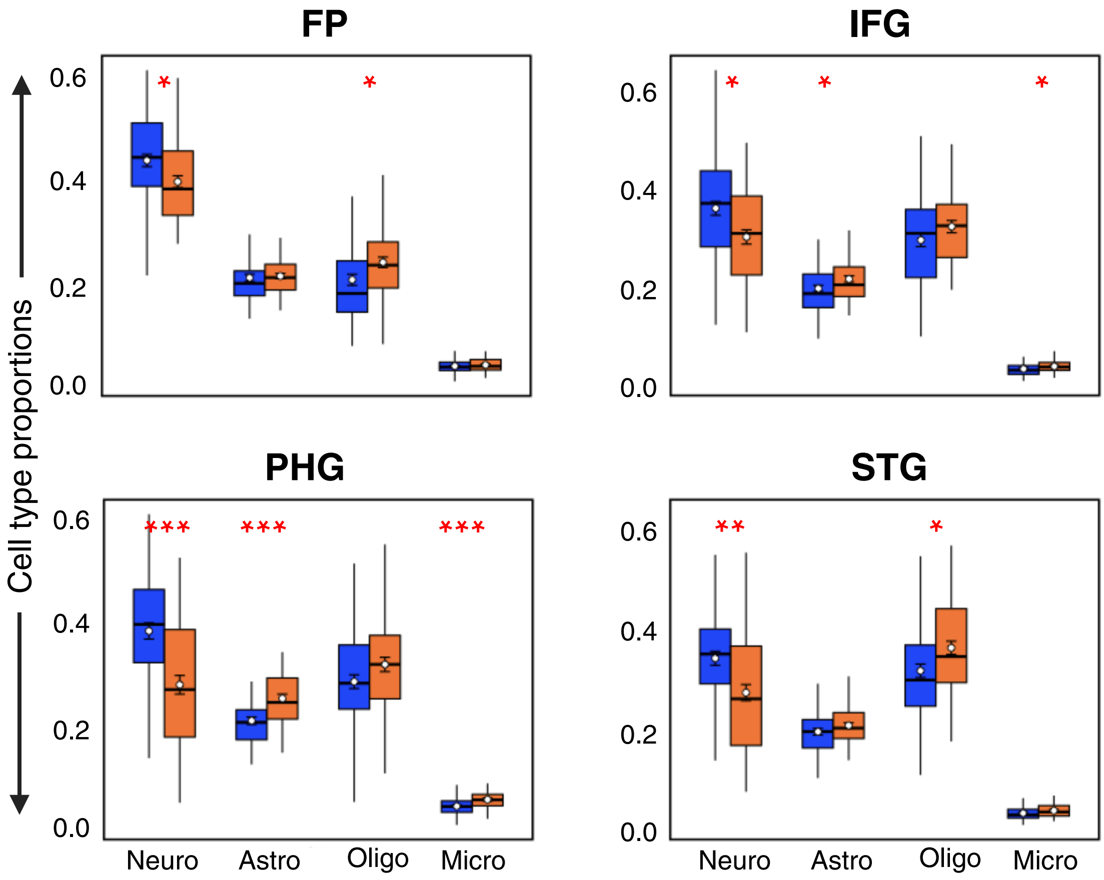


**Fig. S2: Comparative analysis of cell-type proportions across four brain regions using matched samples from 108 individuals.** Boxplots display cell-type proportion estimates across four brain regions, inferred using *dtangle*. Same format, notations and conventions as in Fig. 2A in the main text is used here.

### Supplementary Figure 3

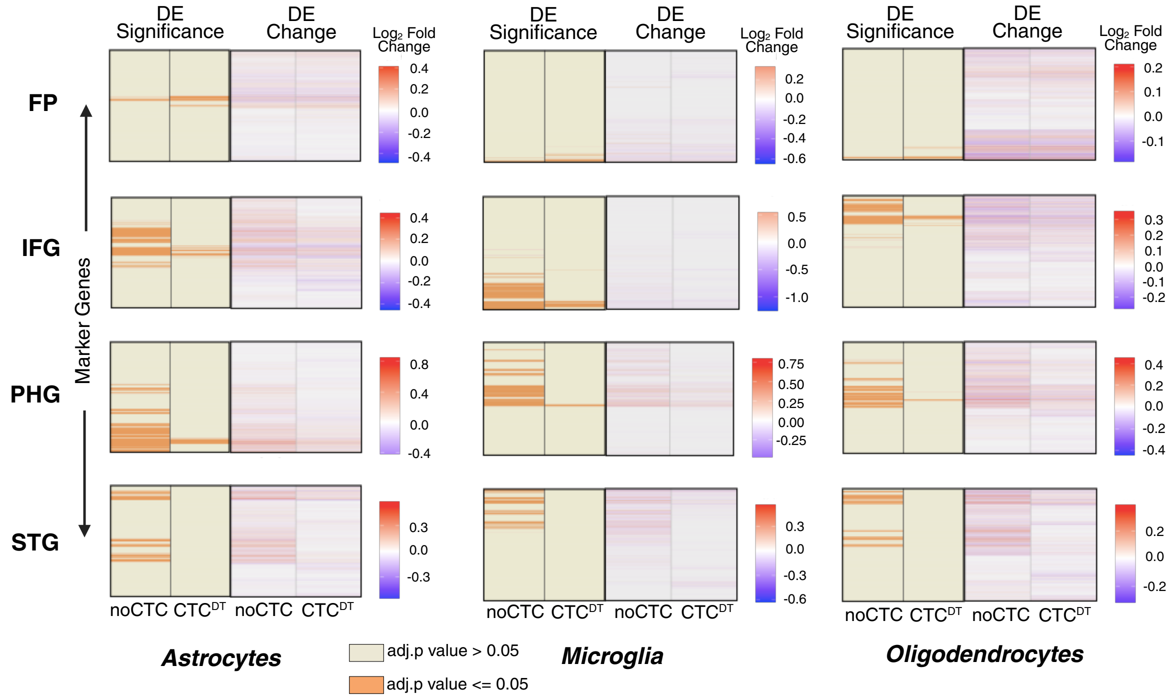


**Fig. S3: Marker gene enrichment across brain regions.** Heatmaps show enrichment patterns of glial marker genes (Astrocyte, Microglia and Oligodendrocyte) derived from the MultiBrain signature matrix for each brain region. In each panel, the left heatmap represents uncorrected analysis (noCTC), while the right heatmap shows results after correction ($CTC^{DT}$). Rows correspond to individual marker genes. The left panels ("DE Significance") show gene-specific adjusted p-values (adj.pvalue), with orange indicating adj.pvalue $\leq 0.05$. The same format, notation, and color scheme as in Fig. 2B of the main text were used here.

**Supplementary Fig. 4**

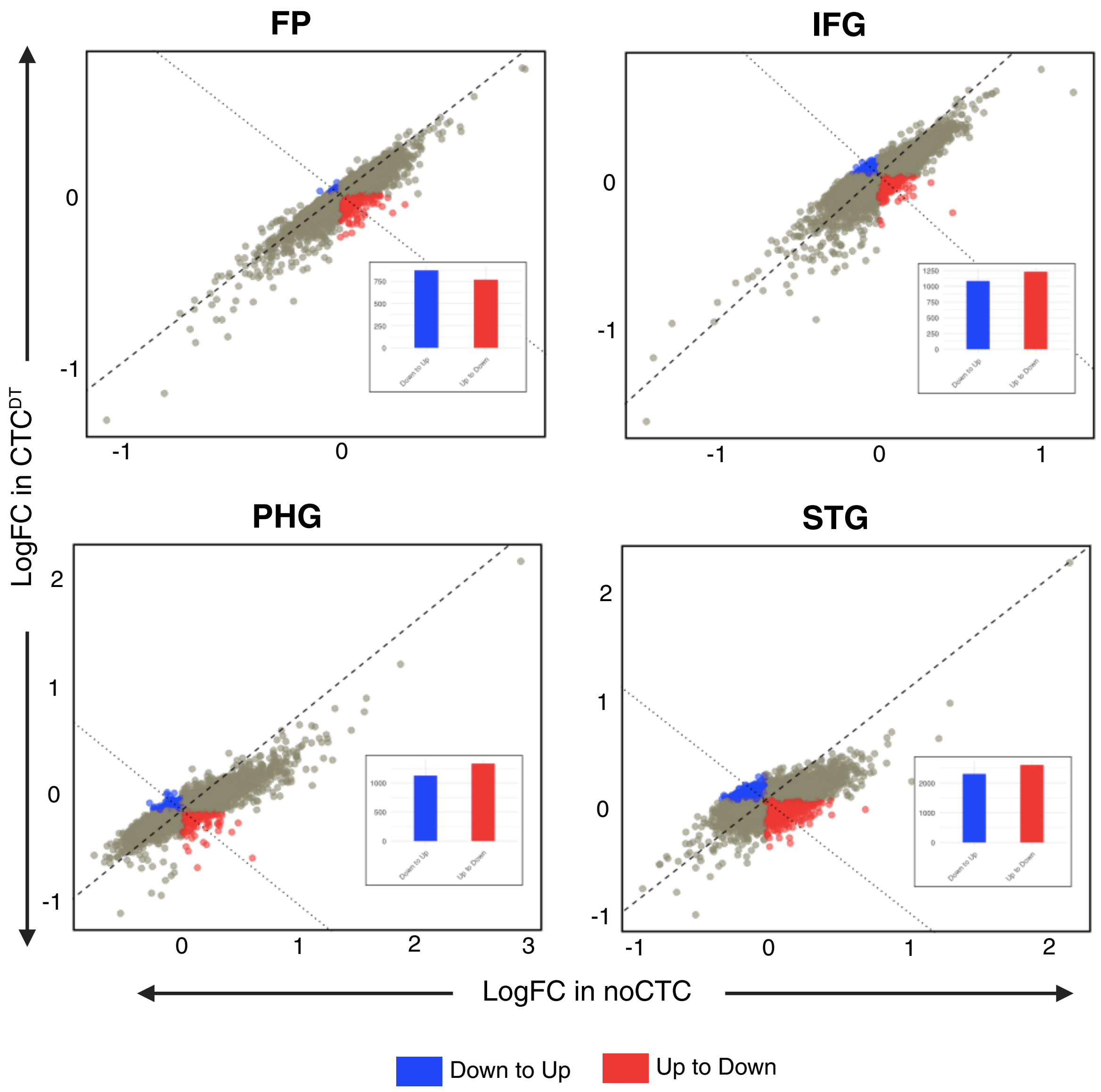


**Fig. S4: Scatter plots show gene-level LogFC across four brain regions, comparing noCTC (x-axis) vs. CTC$^{DT}$ (y-axis).** Points are colored by directional shift: blue (Down to Up), red (Up to Down) and gray (Concordant). Insets summarize the count of genes showing directional reversal (red and blue) for each region.

**Supplementary Fig. 5**

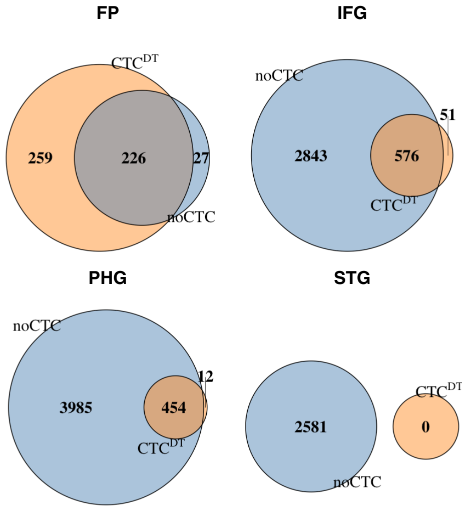


**Fig. S5: Venn diagrams illustrate the overlap of DEGs between noCTC and CTC$^{DT}$ at an FDR threshold of 5%.** The number of shared DEGs between noCTC and CTC$^{DT}$ was 226 in FP, 576 in IFG, 454 in PHG, and 0 in STG. The statistical significance of these overlaps, assessed using a hypergeometric test, yielded the following p values: FP, ~0; IFG, $2.34 \times 10^{-320}$; PHG, $5.89 \times 10^{-229}$; and STG, 1 (no overlap). See also Table 2.

**Supplementary Fig. 6**

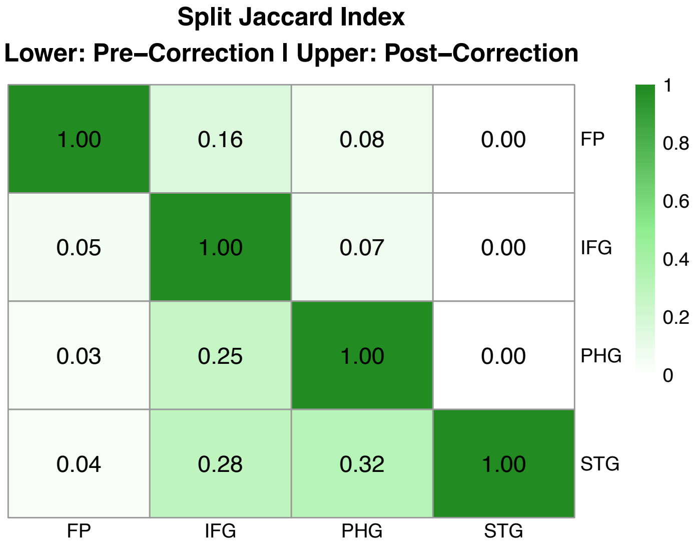


**Fig. S6: Inter-regional transcriptomic relationships across uncorrected and cell-type-aware analyses.** Jaccard similarity indices between brain regions, quantify the extent of overlap in differentially expressed gene (DEG) profiles. Higher similarity values indicate greater convergence in transcriptional signatures between two regions, while lower values reflect region-specific divergence. This analysis captures the heterogeneity of AD-associated transcriptomic remodelling across cortical regions. Details described in Supplementary File 1.

**Supplementary Fig. 7**

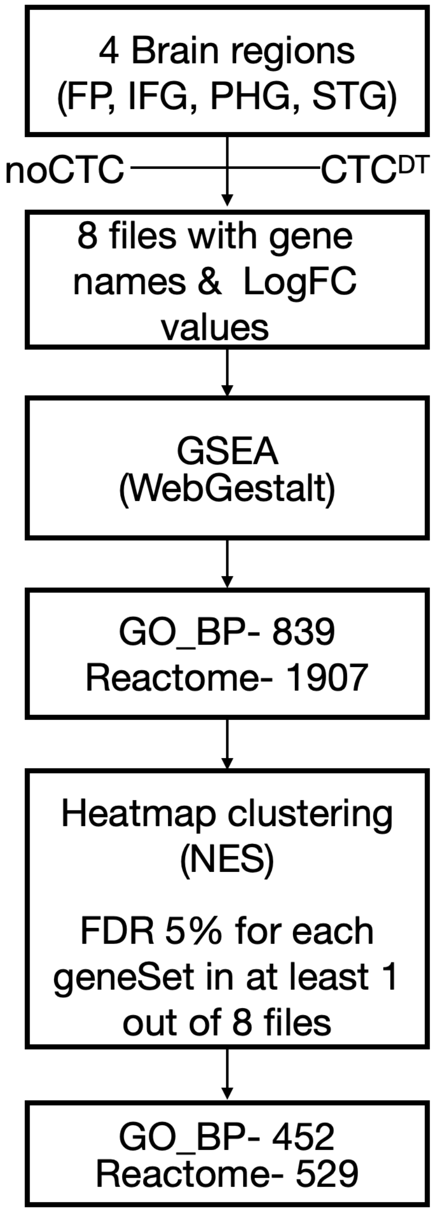


**Fig. S7: Computational workflow for functional enrichment of transcriptional changes in Alzheimer's disease.** Schematic overview of the functional enrichment pipeline applied to differentially expressed genes (DEGs) identified before and after cell-type correction. The workflow includes region-specific DEG detection followed by LogFC-based gene ranking and threshold-free Gene Set Enrichment Analysis (GSEA) using Gene Ontology Biological Process (GO_BP) and Reactome pathway databases. This rank-based approach avoids arbitrary significance cutoffs, thereby aiding pathway-level interpretations of the DE results.

A total of 839 GOBP gene sets and 1,907 Reactome pathways were tested in the initial GSEA analysis. Applying a significance threshold of FDR < 5% identified 452 GOBP terms and 529 Reactome pathways that were significantly enriched for DEGs in at least one of the eight region-specific analyses. These represent the union of pathways meeting the FDR criterion across all regions.

The resulting GSEA profiles were clustered and visualized as heatmaps in Fig. 4.

**Supplementary Fig. 8**

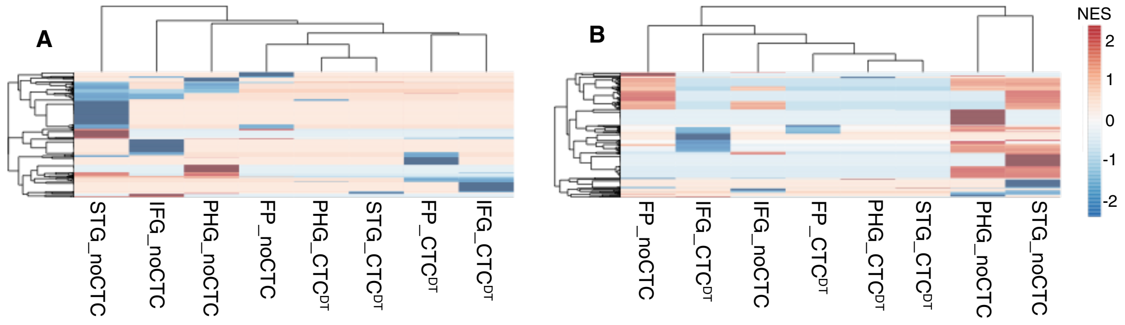


**Fig. S8: Exclusive functional enrichment across brain regions before and after cell-type correction.** Side-by-side heatmaps display enrichment scores for Reactome pathways (A) and GO_BP (B) across brain regions under uncorrected and corrected conditions. Only functions exclusively enriched in either noCTC or $CTC^{DT}$ were retained, highlighting transcriptional shifts specific to each analytical framework. We follow the same notations and conventions as in Fig 4.

Pathway enrichment analysis yielded 46, 111, 100, and 226 exclusively enriched for noCTC data in FP, IFG, PHG, and STG, respectively, compared to 60, 57, 8, and 9 in $CTC^{DT}$. When aggregated across regions, this disparity culminated in 174 pathways exclusive to noCTC and only 58 exclusive to $CTC^{DT}$. A similar pattern as with Reactome pathways was observed for GO_BP enrichment. Across all the regions, 196 GO_BP pathways were uniquely enriched in noCTC, while only 29 were exclusive to $CTC^{DT}$. Region-wise, noCTC yielded the following pathway counts: FP (138), IFG (70), PHG (215), and STG (282), in contrast to markedly lower counts of $CTC^{DT}$- FP (30), IFG (88), PHG (7), and STG (2), except for IFG.

**Supplementary Fig. 9**

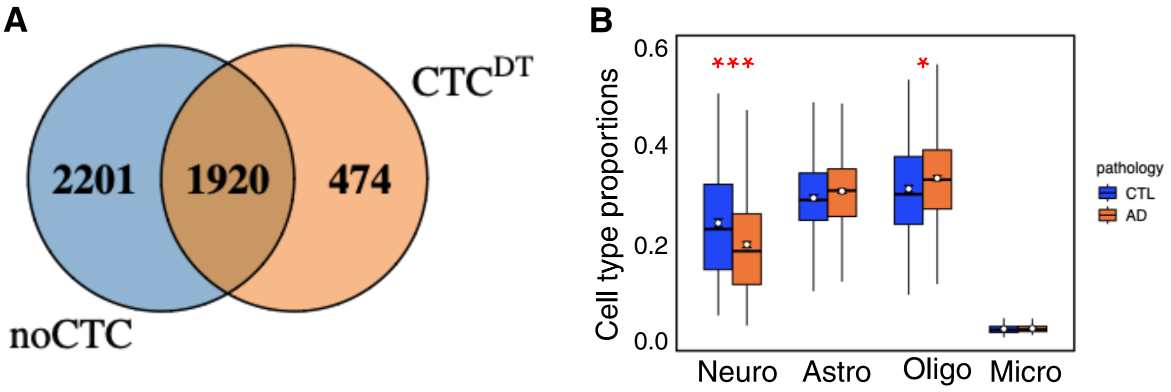


**Fig. S9: Cell-type proportion estimates and DEG overlap in ROSMAP-DLPFC.** (A) Venn diagram illustrates the overlap of differentially expressed genes (DEGs) identified under uncorrected (noCTC) and cell-type-aware corrected ($CTC^{DT}$) models at an FDR threshold of 5%. The shared and unique gene sets reflect the impact of correction on transcriptional signal refinement. (B) Boxplots show cell-type proportion estimates for DLPFC samples, inferred using dtangle. Blue and orange bars represent distinct experimental conditions (e.g., Control vs. AD). Asterisks indicate statistically significant differences (adjusted $p \leq 0.05$ [*], $p \leq 0.01$ *[**], and* $p \leq 0.001$ *[***])*, highlighting compositional shifts associated with disease status.
